# Intrinsic molecular identifiers enable robust molecular counting in single-cell sequencing

**DOI:** 10.1101/2024.10.04.616561

**Authors:** Kristina M. Fontanez, Yigal Agam, Sophia Bevans, Aaron A. May-Zhang, Corey Hayford, Yi Xue, Jacob S. A. Ishibashi, Rachael Komuhendo, Lucas Yoder, Pabodha Hettige, Hannah D. Rickner, Jesse Q. Zhang, Ahmad Osman, Chris D’Amato, Trinity Smithers, Sarah Calkins, Muntasir Rahman, Kiryakos S. Mutafopulos, Sepehr Kiani, Robert H. Meltzer

## Abstract

Particle-templated instant partition sequencing (PIPseq), is an emerging approach for massively scalable single-cell gene expression studies that does not require complex instrumentation or expensive consumables. We present PIPseq^TM^ V, a novel implementation of the PIPseq workflow with significant improvements in assay performance and sensitivity compared to prior methods. Among the innovations driving the improved performance in PIPseq V is a new approach for transcript counting using Intrinsic Molecular Identifiers (IMIs) from the captured transcript sequence, eliminating the need for traditional Unique Molecular Identifiers (UMIs). IMIs are the starting positions for molecules generated by random fragmentation after limited cycle PCR, which can be used to uniquely identify individual transcript copies of genes expressed within individual cells. To correct for experimental variation in IMI generation, we have developed a dynamic correction strategy that can adapt to different sample cell input, transcript expression, and sequencing depth without the need for external benchmarking or controls. Our results demonstrate that PIPseq V with IMI-based analysis provides biological information comparable to established UMI-based approaches while avoiding UMI-associated biases. Dynamic correction offers a robust and data-driven analysis strategy for scRNAseq.

## INTRODUCTION

Single-cell RNA sequencing (scRNAseq) provides unprecedented insights into the biology of the complex cellular milieu, as evidenced by international initiatives like the Human Cell Atlas relying upon these methods to build comprehensive reference maps of all human cells^1^. The rapid growth of single-cell analysis tools offers researchers a diverse range of options. These approaches span from low-throughput plate-based methods ^2,3^ to *in-situ* combinatorial barcoding methods ^4–7^ to high-throughput, bead-based methods using droplets ^8–11^ or wells ^12,13^.

Particle-templated Instant Partitioned sequencing (PIPseq)^11^ is a highly scalable, flexible, and user-friendly method with several advantages that are particularly enabling for single-cell studies. PIPseq eliminates complex and expensive instrumentation and microfluidic consumables, limiting the accessibility of existing single-cell approaches. The emulsification process occurs throughout the entire sample simultaneously, reducing the potential for sample degradation during the sequential droplet creation and cell capture process common in microfluidic methods. PIPseq is highly scalable, with single-cell capture demonstrated from tens of cells to millions of cells in a single-tube format ^11^. The versatility of PIPseq has been shown in a wide range of applications ^11,14–17^, with analytical workflows that have evolved to support the unique biochemistries and experimental modalities enabled by PIPseq.

Here we present PIPseq V, a significant improvement to the previously released PIPseq v4.0PLUS chemistry and workflow. As a recent meta-analysis of scRNA technologies has revealed, there is growing demand for improvements to single-cell analysis accessibility, scalability, sensitivity, cell capture efficiency, and cell type compatibility. ^18^ PIPseq V combines new methods for cell lysis and capture, nucleic acid conversion, molecular counting, and sequencing library preparation that address the limitations of previously published prototype and commercially available PIPseq chemistries ^11^. In combination, these innovations contribute to PIPseq V’s significantly improved performance across a variety of sample types.

## RESULTS

### PIPseq V workflow

The PIPseq V 3’ scRNAseq workflow relies on Particle-templated Instant Partitioning (Figure 1A) to instantaneously capture hundreds to millions of cells (or nuclei) inside microdroplets upon vortexing ^11^. After partitioning, the single-cell emulsion is mixed with a second, non-templated emulsion containing a lysis reagent that is transferred into the single-cell droplets via micellar molecular transport ^19^ (Figure 1B). This is a new method for lysis of captured cells distinct from the heat-activated enzymatic lysis previously described ^11^. The combined emulsion is then incubated at a sequence of specific temperatures to facilitate cell (or nuclei) lysis and RNA capture onto the poly(dT)V capture site. After RNA capture, the emulsion is broken by chemical disruption which releases the core particles with hybridized mRNA (Figure 1C). The particles are washed into a novel reverse transcription reaction that significantly improves reverse transcription and template switching efficiencies, contributing to improved gene and transcript sensitivity in PIPseq V. The result of reverse transcription is a barcoded complementary DNA (cDNA) copy of each hybridized mRNA with a PCR handle on the 3′ end (Figure 1D). Next, the core particles containing cDNA are washed into a PCR amplification buffer with primers complementary to the PCR handles provided by the core particle and TSOs. Importantly, PCR amplification is limited to a fixed number of cycles which sets a ceiling on the number of possible cDNA copies in the supernatant (Figure 1E). cDNA amplification cycles are fixed, regardless of sample input, cell type, or kit scale. This is another distinct change from previous PIPseq workflows which recommended different amplification cycle numbers for different cell input and sample types. The resulting amplified cDNA product is then isolated from the core template particles and randomly fragmented by enzymatic digestion with DNA endonuclease (Figure 1F). This random fragmentation step results in unique cut sites that can serve as Intrinsic Molecular Identifiers (IMIs) which are the cornerstone of the novel molecular counting approach implemented in PIPseq V. All of the DNA fragments are then processed through end-repair, A-tailing, ligation, and amplification of indexed adapters appropriate for next-generation sequencing (Figure 1F). Subsequent data analysis is performed using PIPseeker software as described in detail in Methods.

**Figure 1.**
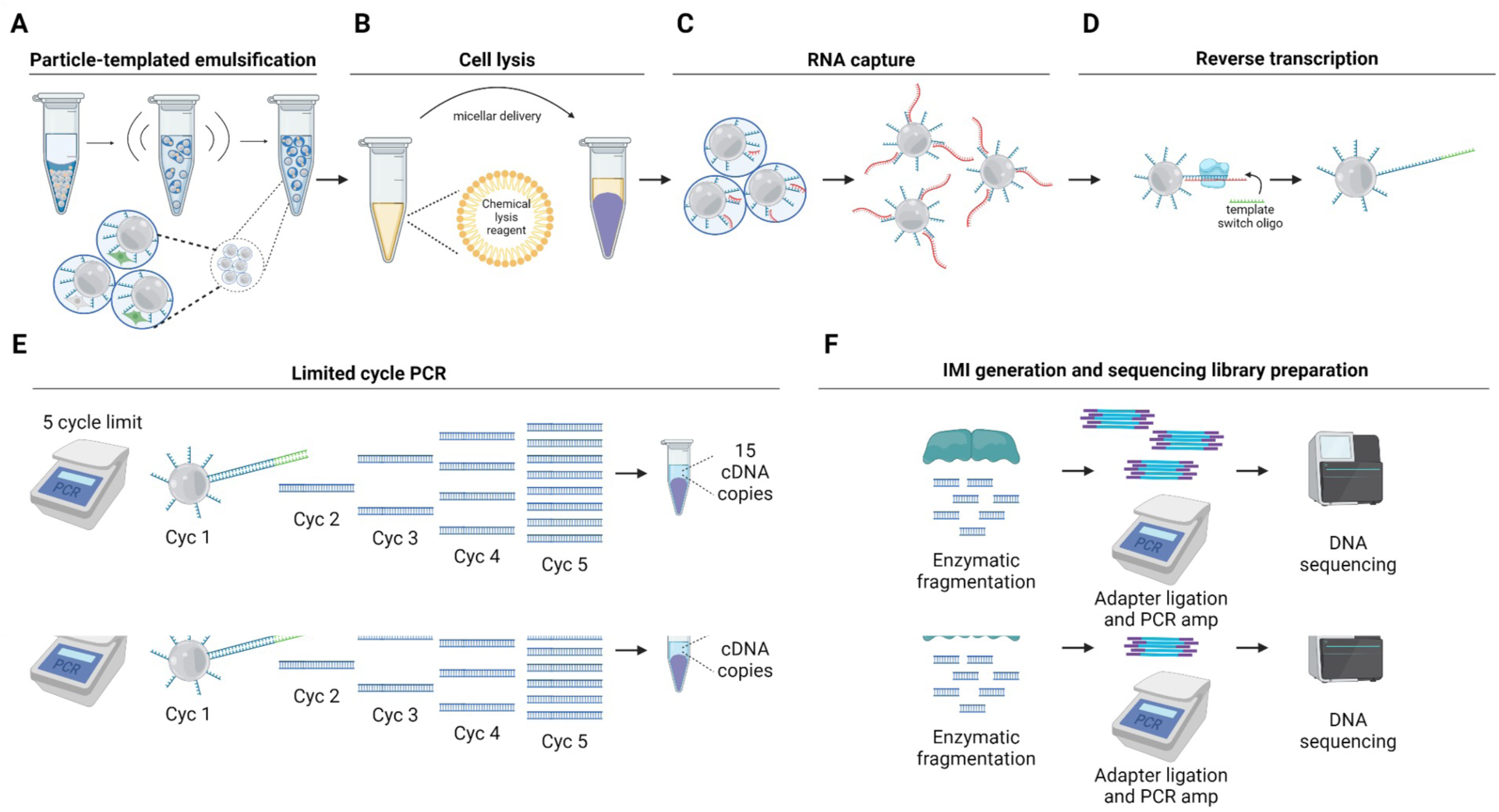
PIPseq V workflow. A) Barcoded hydrogel beads are combined with cells/nuclei and Partitioning Reagent, then vortexed at 3,000 rpm for 135/140 sec with PIPseq Vortex Mixer to generate a PIP emulsion. B) Chemical lysis reagent emulsion is added to the PIP emulsion and incubated in PIPseq Dry Bath to facilitate cellular/nuclear lysis in the droplets. C) After cellular/nuclei lysis in the emulsion, the released mRNA will be captured by the barcoded Poly(dT)V capture moiety on the hydrogel beads. D) PIP emulsions are broken and isolated hydrogel beads carrying mRNA are washes are performed to remove residual impurities. Captured mRNA on hydrogel beads undergo mRNA reverse transcription (RT) to synthesize cDNA. E) Following cDNA synthesis, a limited 5 cycle PCR occurs to generate 15 copies of barcoded cDNA. F) Amplified cDNA supernatant is isolated from the beads and purified. Enzymatic fragmentation on isolated cDNA generates the IMI and sequencing ready libraries are prepared on these fragments for Illumina sequencing.

### PIPseq V is compatible with a diversity of sample types

PIPseq V performance was evaluated across multiple cell types with varying intrinsic mRNA content in kits scaling from 2,000 to 100,000 targeted cell capture. Model sample types included mixed mouse and human cultured cells, human peripheral blood mononuclear cells (PBMCs), and nuclei prepared from liquid nitrogen frozen mouse brian. Nuclei preparations were also evaluated with and without sample fixation. This panel of test samples demonstrates the scalability of the PIPseq platform, compatibility across species and tissues, and utility across simple to very complex tissues.

### Mixed cultured cells

Human HEK 293T and mouse NIH 3T3 cell line mixtures are a common model system for single-cell benchmarking because of their consistent performance in single-cell assays as they are high transcript-expressing cells and enable species mixing experiments to determine multiplet rates ^9^. HEK / 3T3 cell mixtures were prepared as described in the Methods and 5000 mixed cells were loaded to PIPseq V T2 (Figure 2). 3,611 cells were detected in PIPseeker analysis, resulting in >72% cell capture with 9,437 median transcripts per cell and 3,415 medium genes per cell at 20,000 RPCC (Table 1). The barcode rank plot is representative of high sample quality (Figure 2A), while UMAP and barnyard plots display high resolution of species-specific cell clusters (Figure 2 B-D). The observed fraction of species mixing is less than 5%, and this low multiplet occurrence contributes to the increased sequencing efficiency and sensitivities with PIPseq V (Figure 2D).

**Figure 2.**
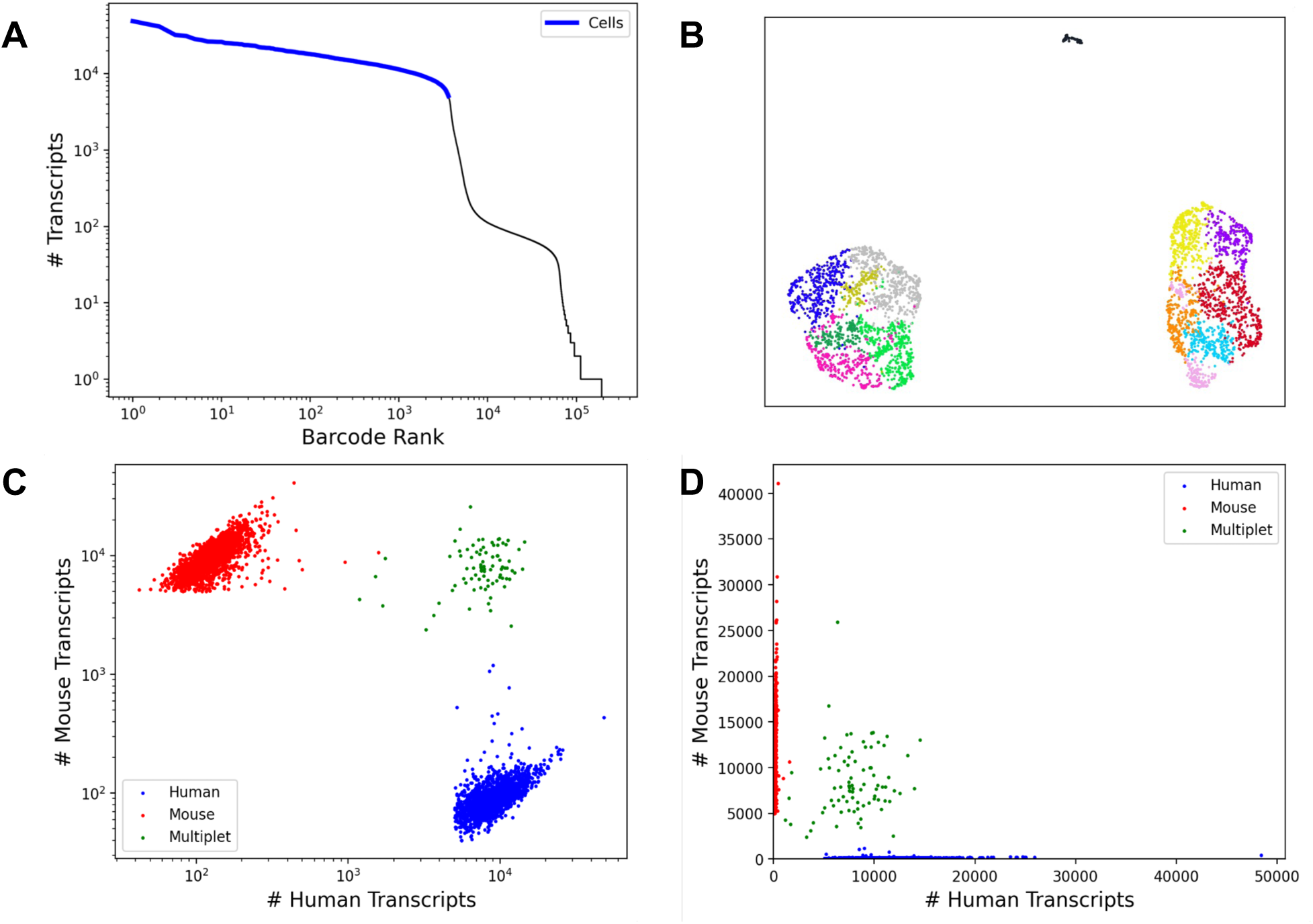
Mouse / human cell separation in PIPseq V. 4,106 HEK 293T/NIH 3T3 cells captured in PIPseq V T2. A) Barcode rank plot of high quality sample recovery. B-D) UMAP and ‘barnyard’ scatter plots demonstrating clean mouse and human species separation and low multiplet rate (<5%).

**Table 1.** PIPseq V chemistry efficacy across a range of sample type and kit sizes, including DSP-methanol fixed mouse brain nuclei.

| 5 Cycle Data<br>Sample Type | Kit size | Sequencing Depth<br>(Reads Per Cell) | Median<br>Transcripts<br>Per Cell<br>(IMI) | Median<br>Genes Per<br>Cell (IMI) |
| --- | --- | --- | --- | --- |
| HEK/3T3 | T2 | 20,000 | 9,437 | 3,415 |
| PBMC | T20 | 20,000 | 4,402 | 1,871 |
| Mouse Brain<br>Nuclei (Unfixed) | T100 | 17,068 | 4,532 | 2,166 |
| Mouse Brain<br>Nuclei (Fixed) | T20 | 17,000 | 2,988 | 1,914 |

### PBMCs

The PIPseq V workflow was demonstrated in peripheral blood mononuclear cells (PBMC) by loading 40,000 cells into the PIPseq V T20 3’ Single Cell RNA assay and analyzed through PIPseeker. High cell type diversity and variable transcript expression levels in PMBCs make them ideal for demonstrating the sensitivity of PIPseq V performance. 31,613 cells were identified by the PIPseeker analysis (75% capture efficiency) and demonstrated clear separation from non-cell containing background barcode (Figure 3A) The identified PBMCs yielded 4,402 median transcripts per cell and 1,871 median genes per cell at 20,000 RPCC. Automated cell type annotation identified a robust range of immune cell populations including large CD14 monocyte and B cell clusters (Figure 3B-C). Rare HSPC and cytotoxic CD4 T cells were also identified. Canonical marker gene recovery such as CD74 in B cells, GNLY in natural killer cells, and CCR7 in naive CD4+ T cells, highlights PIPseq V identification of biological differentiation among cell clusters (Figure 3D).

**Figure 3.**
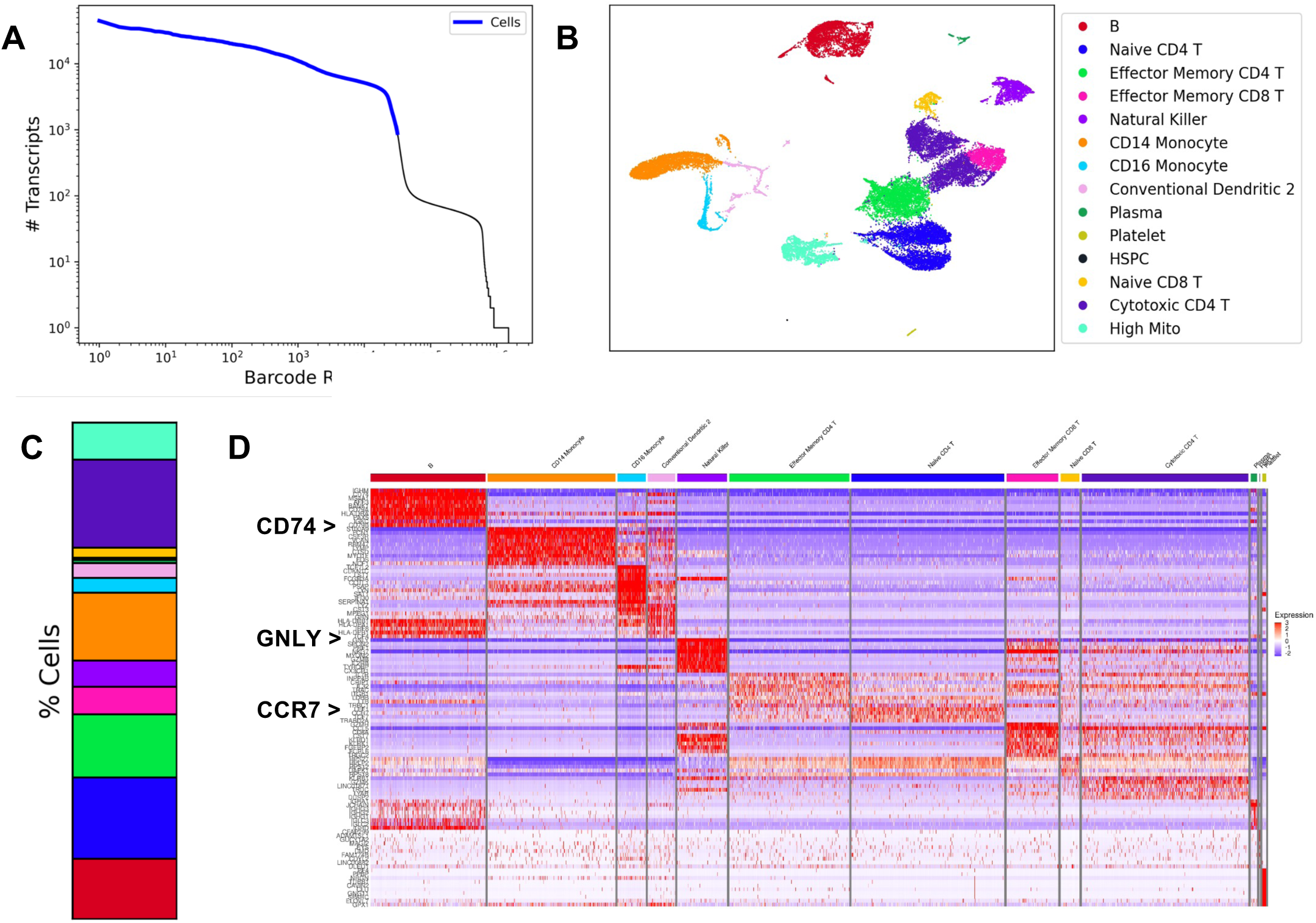
PBMCs resolved by PIPseq V. 40,000 human PBMCs loaded into PIPseq with 31,613 cells identified. A) Barcode rank plot demonstrating high quality sample capture. B) Cell type annotated UMAP with clear immune cell cluster separation and identification. C) Percent composition of PBMC cell type clusters. D) Heatmap representation of the top 10 expressed genes per cell type cluster, including canonical markers such as CD74 (B cells), GNLY (natural killer cells), and CCR7 (naive CD4+ T cells).

### Mouse brain nuclei

Finally, PIPseq V compatibility with both DSP-methanol fixed and unfixed mouse brain nuclei samples was validated. Nuclei extracted from frozen mouse brains are a highly diverse, low mRNA-containing sample type. High sample input is necessary to identify the spectrum of interacting neuronal and glial populations in these complex tissues. From 200,000 cells loaded, 155,000 unfixed mouse brain nuclei were captured (77.5% capture efficiency) with the PIPseq V T100 3’ Single Cell RNA Kit (Figure 4), with 4,532 median transcripts per cell and 2,166 median genes per cell at 17,068 RPCC (Table 1). In parallel, isolated nuclei were fixed with a DSP-Methanol solution, and 40,000 nuclei were loaded to PIPseq V T20 (Figure 4). 34,679 nuclei were identified by PIPseeker analysis (86.7% capture efficiency) with 2,988 median transcripts per cell and 1,914 median genes per cell recovered at 17,000 RPCC (Table 1). Both fixed and unfixed mouse brain nuclei samples produced well-defined UMAP clustering and automated cell type annotations, with a wide variety of brain cell populations represented (Figure 4A-B). Other than an increase in vascular cell recovery with DSP-methanol fixation, there are minimal differences in cell type representation between the fixed and unfixed sample types (Figure 4A-C). Notably, we observed multiple glial cell populations, including astrocytes, oligodendrocytes, and microglia, with PIPseq V in both T20 and T100 kit scales (Figure 4A). These challenging cell populations are notably difficult to recover in microfluidic-based single-cell workflows and are implicated in a wide range of neurological diseases ^16^.

**Figure 4.**
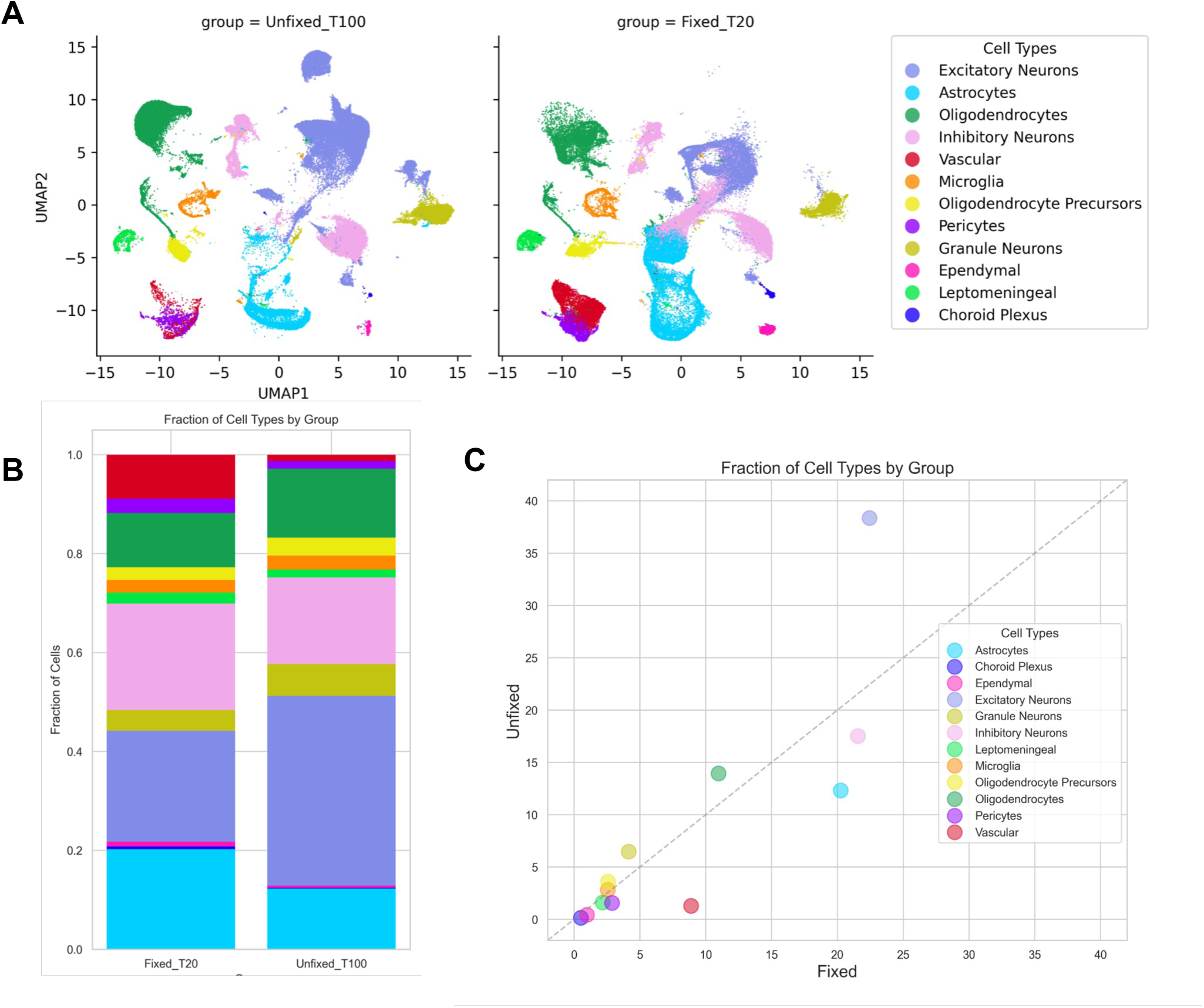
Mouse brain nuclei resolved in PIPseq V. PIPseq V T100 3’ Single Cell RNA Kit captured 155,000 unfixed mouse brain nuclei captured and PIPseq V T20 3’ Single Cell RNA captured 34,679 DSP-methanol fixed mouse brain nuclei. A) UMAP demonstrating high cell type resolution that is comparable between fixed and unfixed samples. Percent composition B) and fraction of cell types by group C) of cell types between the fixed and unfixed samples.

### PIPseq V is optimized for gene and transcript sensitivity

PIPseq V incorporates improvements to cell lysis and capture, nucleic acid conversion, molecular counting, and sequencing library preparation. We compared the result of these changes incorporated against the previously released PIPseq v4.0PLUS platform. 40,000 Human PBMCs were input into the PIPseq v4.0PLUS and PIPseq V kits which yielded 27,220 cells (68% capture) and 31,613 cells (79% capture), respectively (Figure 5A). The datasets were merged and batch-corrected with Seurat integration, resulting in a combined UMAP, with the majority of identified cell populations overlapping (Figure 5A). Violin plots of gene and transcript sensitivity reveal that the PIPseq V workflow results in higher sensitivity across cell types than the PIPseq v4.0PLUS workflow overall (Figure 5B-C). Among all differentially expressed genes (abs. log_2_FC>=0.001), only 0.1% of genes were significantly (p<0.05) differentially expressed with meaningful fold changes (abs. log_2_FC >= 1.5). About one-third of these genes were mitochondrial or ribosomal in origin, indicating that differential expression patterns were similar across workflows (Figure 5D). The average expression within PBMC cell types for the intersection of all expressed genes (p<0.05) was highly correlated (R>=0.96 for all cell types) for the PIPseq V and PIPseq v4.0PLUS workflows (Figure 5E). Observed expression for most genes was higher in the PIPseq V workflow, as indicated by the genes above the dotted line (Figure 5E).

**Figure 5.**
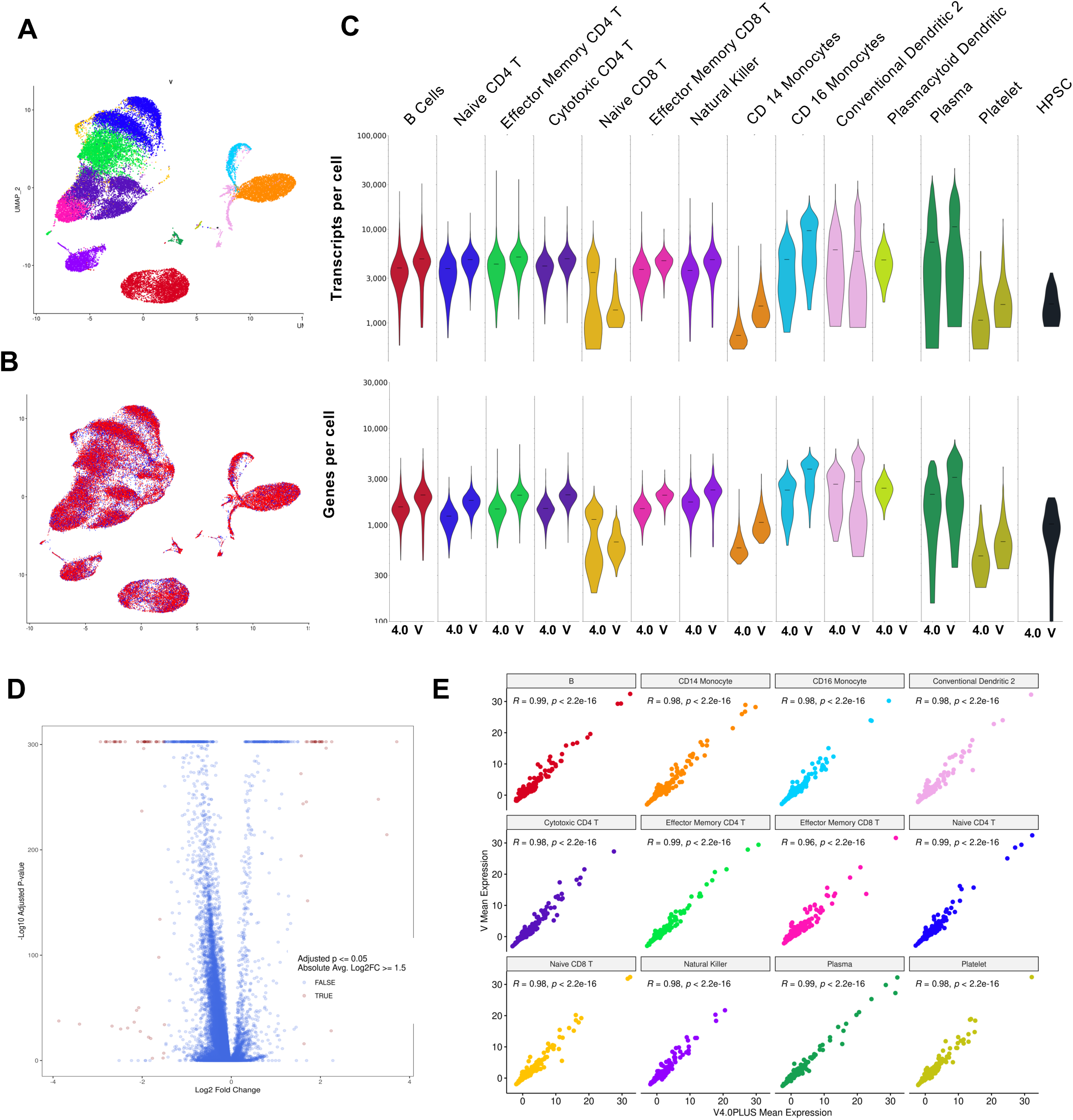
PIPseq v4.0PLUS vs PIPseq V. 40,000 PBMCs were processed with PIPseq v4.0PLUS and PIPseq V. A) PIPseq V analysis resolves diverse immune cell populations. Color coding is maintained across figure panels. B) Merged and batch corrected UMAP of PIPseq V (red) and PIPseq v4.0PLUS (blue) datasets indicate significant overlap in identified PBMC cell type populations. Violin plots comparing v4.0PLUS and V workflows transcript/cell (B) and genes/cell (C) distributions across different cell types identified in PBMCs show that the PIPseq V workflow yields higher overall sensitivity across different cell types. D) Only 0.1% of all genes are significantly differentially expressed between the PIPseq v4.0PLUS and PIPseq V workflows. Positive = higher expression in V workflow, negative=higher expression in the V4.0PLUS workflow. E) Gene expression in PBMC cell types (avg >1^−15^) is highly correlated (R>=0.96 overall) between the two workflows.

### Intrinsic molecular identification - new methods for transcript counting

The most significant variation from previous single-cell methods introduced with PIPseq V is the adoption of Intrinsic Molecular Identifiers (IMIs) to correct for PCR-dependent duplication of sequencing reads. Rather than using an extrinsic, randomer tag to identify individual source mRNA molecules, PIPseq V relies on randomized fragmentation of early copy cDNA molecules to provide identifying information. (Figure 6). The transcript sequence thereby encodes information about the gene identity and fragmentation position, which can be used to trace the molecular origin of each sequencing read in a data set.

**Figure 6.**
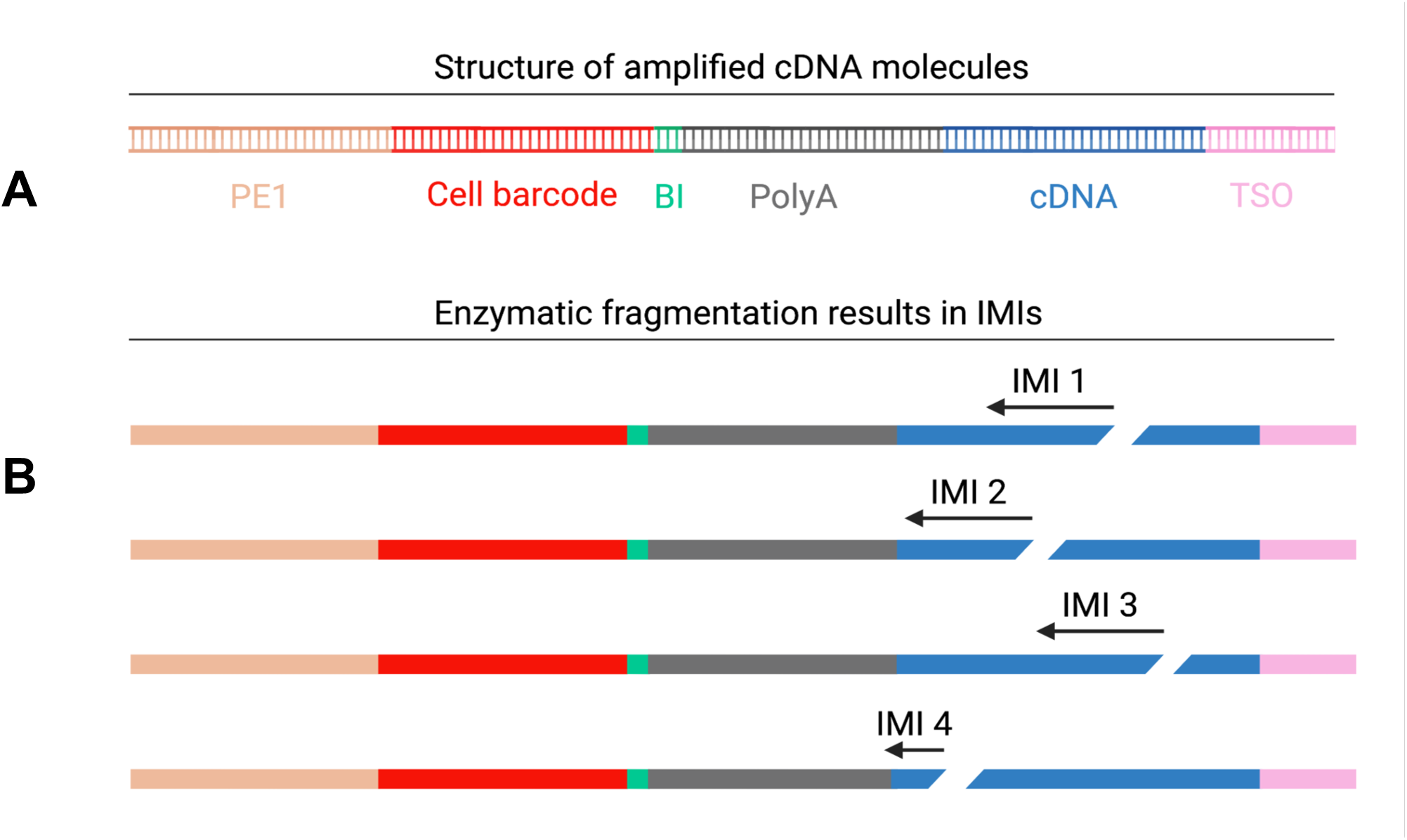
Random enzymatic fragmentation sites create intrinsic molecular identifiers (IMIs). Up to 15 different IMIs can be created from a single captured molecule generated by 5 cycles of PCR amplification.

An important advantage of eliminating poly-N encoded UMIs is the reduced opportunity for mRNA selection bias. For comparative experiments, we manufactured PIPseq beads configured for IMI-based analysis (Figure 7A), or for traditional UMI-based analysis (Figure 7B). With UMI-containing PIPseq particles (UCPs), we observe a poly-T bias towards the end of the UMI (Figure 7C), as has been previously observed in other 3′ scRNAseq technologies ^20,21^. This represents the increased probability of mRNA poly-A tailing preferentially selecting T-enriched sequences in canonical UMI structures that effectively decrease the available UMI space. Use of the intrinsic fragmentation site as the identifying feature for each molecule, in contrast, imposes no selection bias in the identifying sequences. (Figure 7D). For improved molecular counting estimation, we have also introduced a short, 3-base binning index in our bead barcode sequences - by limiting the variable region of this binning index and introducing a fixed spacer sequence between the binning index and the poly-T capture moiety, selection bias is not observed in the binning indices (Figure 7E).

**Figure 7.**
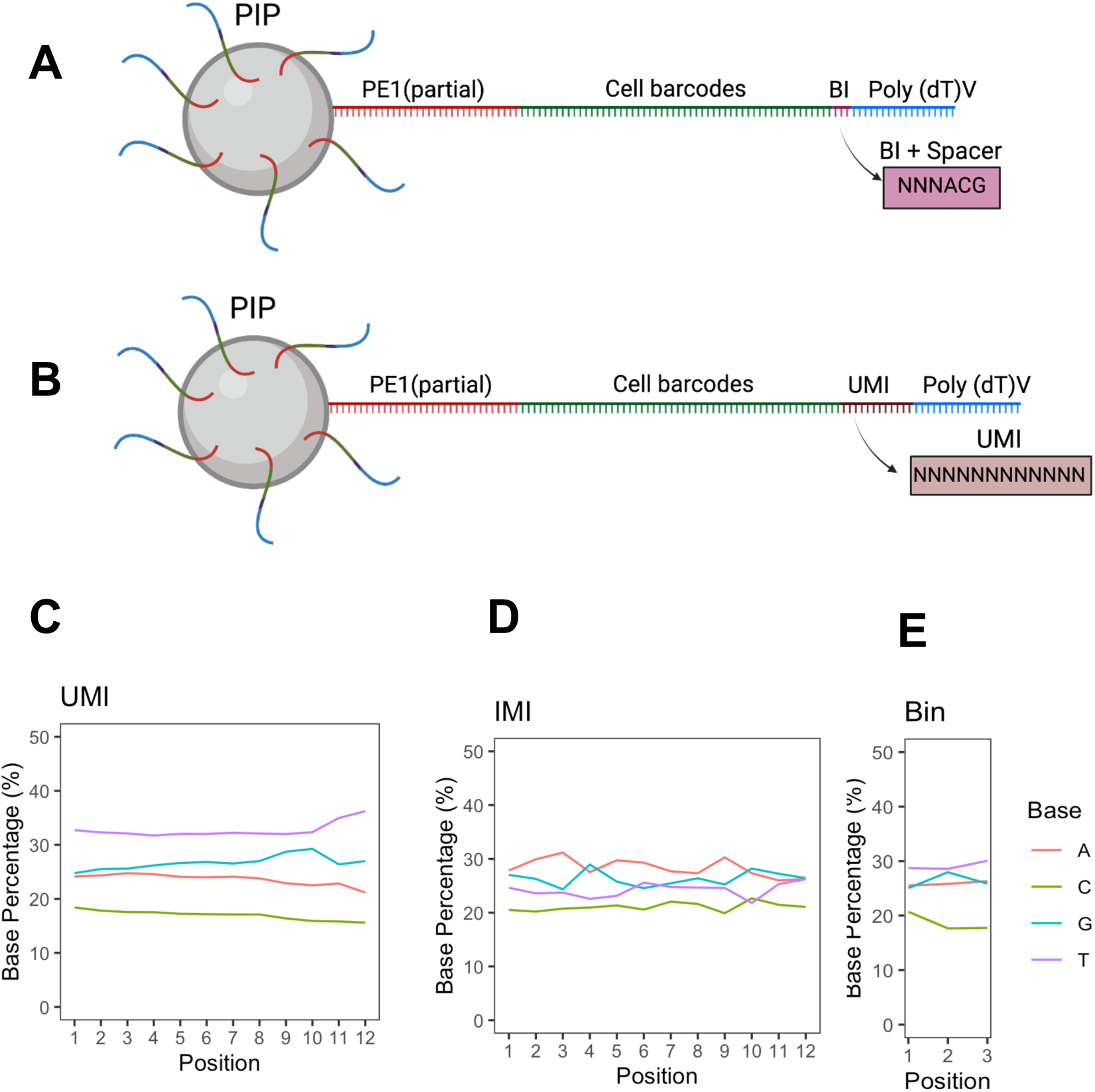
Hydrogel bead architecture facilitating barcoded mRNA capture for IMI analysis. A) PIPseq V core particles with oligonucleotide sequences carrying 5’ PCR handle, cellular barcodes, 3N base binning index, ACG spacer, and poly A mRNA capture moiety. B) UMI-containing core particles (UCP) oligonucleotide sequences carrying 5’ PCR handle, cellular barcodes, 12N base UMI, and poly A mRNA capture moiety. C) Base distribution of unduplicated reads vs. position in UMI when UCPs are used, D) IMI when IMI particles are used, E) Binning index when IMI particles are used. UMIs are enriched for T bases while IMIs and Bin indexes have expected random mixture of bases with less T bias.

UMI-based methods rely on the incorporation of a unique molecular identifier (UMI) before PCR amplification. Under ideal conditions, each PCR-generated daughter molecule inherits the parent molecule’s unique sequence tag, enabling downstream deduplication of PCR copies for accurate quantification of original transcripts. In contrast, the IMI-based workflow relies upon the endogenous generation of IMIs via random enzymatic fragmentation, *after* PCR amplification. The PIPseq V workflow incorporates two critical aspects that enable this strategy, (1) PCR cycle count limitation and (2) enzymatic fragmentation of cDNA following limited cycle PCR amplification. Both steps have a significant impact on the accuracy of this novel molecular counting strategy.

As one double-stranded cDNA molecule remains bound to the PIPseq core template particle, the number of expected amplified cDNA copies from each original mRNA transcript is *x* = (2^*n*−1^) − 1, where *n* is the number of amplification cycles. Ideally, 4 PCR cycles will result in 7 cDNA copies per transcript and 5 cycles will result in 15 copies. This expected replication of the original molecules, however, is countered by inefficiencies through the amplification and library conversion process. Amplicon losses may occur due to a combination of volumetric transfer efficiency, fragmentation of cDNA to an appropriate range of lengths, ligation efficiency, and library amplification efficiency. These dependencies may vary from experiment to experiment, and therefore the total number of amplicons originating from an individual mRNA transcript will exhibit experimental variability.

### Observed molecule conversion efficiency

We used UCPs to perform PIPseq experiments using multiple sample input types for direct comparison of UMI-based and IMI-based molecule counts for quantitative estimation of WTA-based inflation (occurs when more than one IMI is generated from a single original parent molecule). Across all sample types, when evaluating the observed distribution of IMIs per UMI, there are far fewer instances of 15 unique IMIs than would be expected assuming 100% WTA efficiency of a 5-cycle PCR (Figure 8), and in fact, the majority of UMIs had only one associated IMI. These data demonstrate that transcript conversion efficiency varies across sample types, as may be expected for samples of varying intrinsic RNA content. Any attempt to correct molecule counts based upon an assumption of 100% process efficiency must necessarily result in a significant underestimation of the actual number of initial RNA molecules present. For a non-UMI-based molecular counting method to provide useful and reliable estimates of transcripts present in individual cells, it must robustly accommodate sample, sequencing depth, and experimental variation. To this end, we have developed a new algorithmic approach for estimating conversion efficiency (Molecule Conversion Efficiency Estimation algorithm), the basis of our Dynamic IMI Correction strategy.

**Figure 8.**
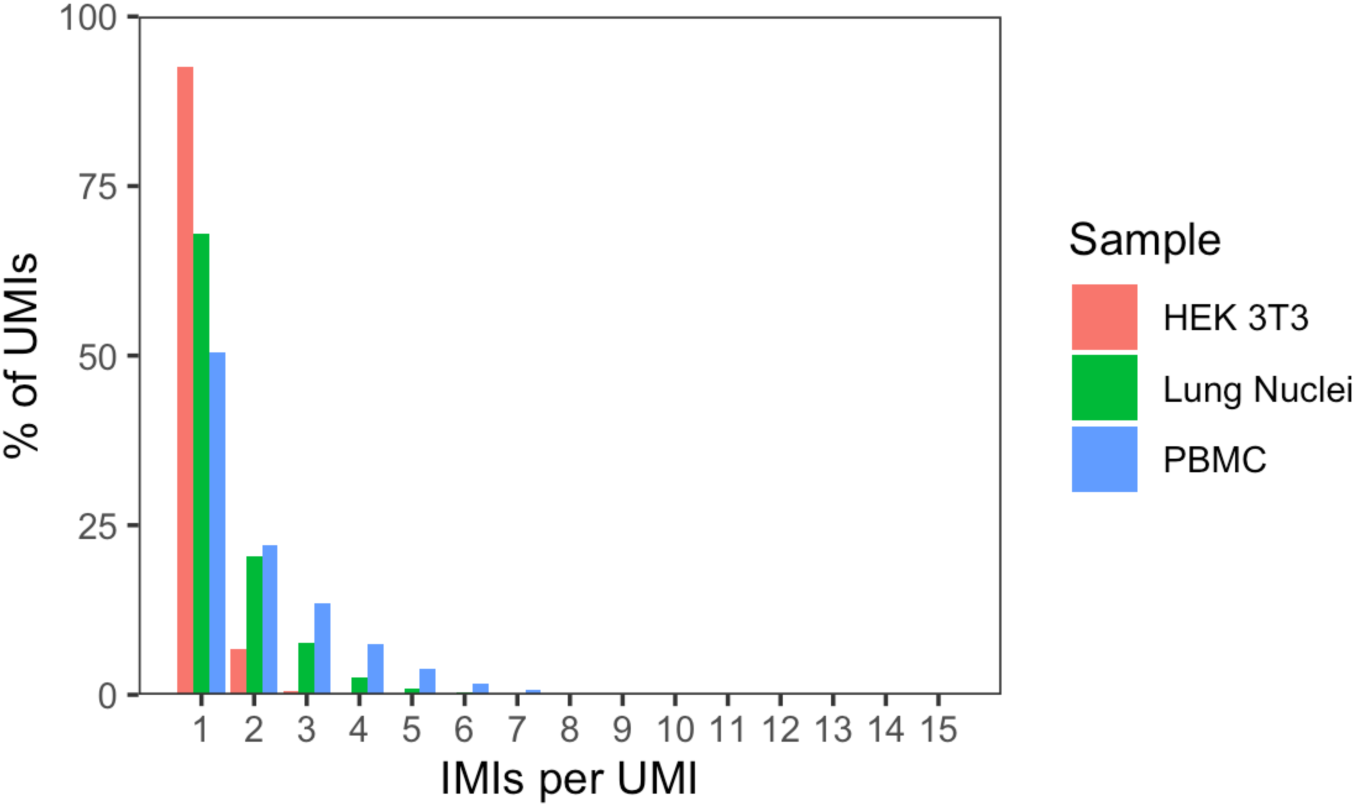
Observed amplification of transcripts counts with 5 cycle IMI analysis. PIPseq experiments were performed in three sample types with UCPs. IMIs observed for each UMI are reported. Optimal process efficiency would result in 15 IMI per UMI across samples.

### Dynamic IMI Correction

The key concept of the Molecule Conversion Efficiency Estimation (MCEE) algorithm relies on the stochasticity of observation of individual reads associated with individual transcripts. By introducing a short, 3-base “binning index” (BI) with the cell barcoding sequences on the PIPseq bead (Figure 7A), we can provide an estimate of the molecule conversion efficiency in any given experiment without the need for spiked-in molecular controls or extensive platform benchmarking. This allows us to dynamically adjust the magnitude of IMI correction according to the degree of IMI inflation in a given dataset.

The analytical workflow (Figure 9) begins the analysis of PIPseq V sequence data by mapping each read against a reference database to identify genes and to enable all reads to be segregated by gene identity. Next, within each cell barcode and gene, individual IMIs are identified without regard for the BI. As the IMI sequences represent intrinsic information encoding the gene identity and fragment starting position as opposed to a randomly assigned barcode sequence, sequencing errors may be corrected by merging IMIs within a Hamming distance of 1. When such similarity is detected, the less frequent IMI is merged into the more frequent IMI. IMIs are then segregated into 64 bins, based on the 3-base BI sequence in each read. Within each bin, the number of unique IMIs is determined by collapsing duplicate IMIs into a single count. This step resolves duplication which occurs during post-fragmentation amplification, and mitigates the probability of observing identical DNA fragmentation sites from distinct transcripts from the same gene.

**Figure 9.**
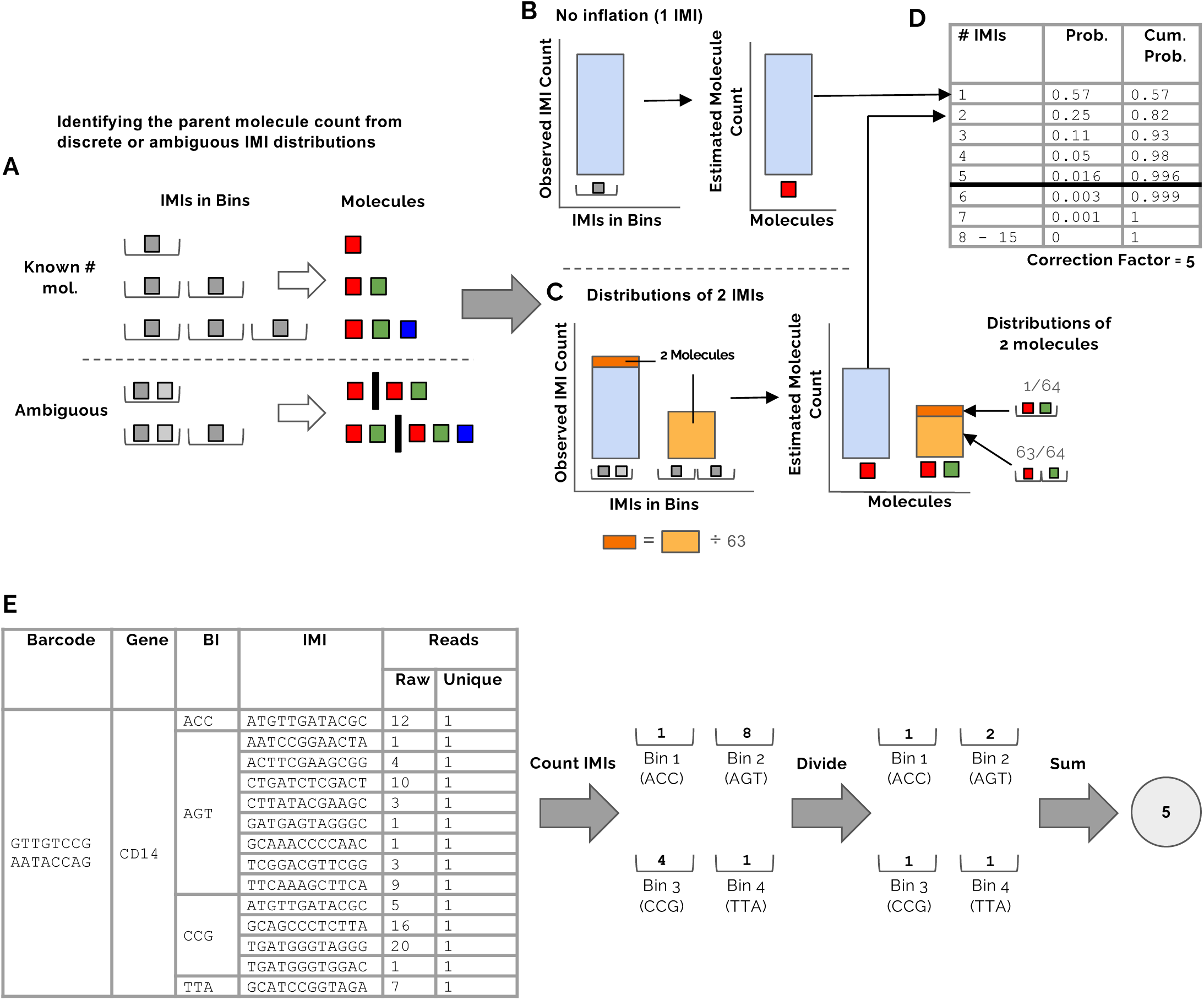
Molecular counting for PIPseq V chemistry. Estimating molecule conversion efficiency and setting a correction factor. A) Examples of known and ambiguous molecular counts for different distributions of IMIs in bins. B-C) Using known molecular counts and combinatorial possibilities of the 3-base binning index to estimate the counts of a single molecule producing different numbers of IMIs. This example illustrates the process for 1 IMI (no inflation, B) and 2 IMIs (C). The count of 1 molecule producing 2 IMIs is corrected by subtracting 1/63 of the count of 2 IMIs in 2 bins, based on the probabilities for the distribution of 2 IMIs (right). Following correction (C, right), the estimated molecule counts (light blue) are used to build a set of conversion probabilities (D). We select a correction factor such that the cumulative conversion probability exceeds 99% (5, in this example). This means that at most, 1% of counts will be inflated. E) Molecular counting. For each barcode-gene combination, IMIs are sorted within bins and collapsed into unique counts. The unique counts are summed and divided by the correction factor. The estimated count for the barcode and gene is the sum of corrected counts from all the bins.

After obtaining counts of unique IMIs in each bin, we estimate molecule conversion efficiency with the MCEE algorithm, and then apply dynamic IMI correction to resolve IMI inflation. The Molecule Conversion Efficiency Estimation algorithm is described in detail in the Methods below (see Analytical methods). We use a subset of the barcode+gene combinations (BGCs) to estimate the conversion distribution, i.e., the probabilities of a single parent molecule producing any number of copies from one to fifteen during WTA. This subset includes all BGCs in which the number of unique IMIs is lower than or equal to fifteen. Some IMI configurations definitively point to a certain number of parent molecules. For example, for a given cell barcode, three unique IMIs observed across three bins can only arise from three unique molecules. In other cases, such as three unique IMIs observed in two bins, the number of original molecules is ambiguous, since multiple IMIs in the same bin can represent distinct molecules or copies of the same molecule fragmented at different locations. This approach assumes that IMIs from the same parent molecule will have the same BGC. For each number of unique IMIs from one to fifteen, the BGCs where the number of molecules can be determined unequivocally are used to estimate the true number of IMIs that can be attributed to conversion from a single parent molecule in a single bin. This is achieved through an iterative process that uses definitive IMI (and parent molecule) configurations as a starting point to probabilistically allocate counts from ambiguous IMI configurations to their constituent parent molecule configurations. By progressively deconvoluting ambiguous counts, we arrive at a refined estimation of single-parent molecule counts within individual bins for each IMI value ranging from 1 to 15. Finally, each of these fifteen estimated counts in single bins is divided by the sum of counts to produce the probability of one molecule producing this many IMIs.

After establishing the conversion probabilities, we perform dynamic IMI correction by using the probabilities to select a correction factor. This is an integer, chosen such that no more than 1% of molecules will be over-counted; the cumulative probability is at least 0.99. Within each BGC, the number of unique IMIs in each bin is divided by the correction factor and rounded up to the nearest integer. The divided counts are then summed across bins to produce the final count for this BGC.

### Evaluating the Molecule Conversion Efficiency Estimation algorithm

The MCEE algorithm estimates the probability of converting a single parent molecule to each possible number of IMIs using only the observed IMIs, binning indexes, and BGCs. The magnitude of IMI inflation (conversion probabilities for >1 IMI) for a given dataset depends on factors such as cell type and sequencing depth. Still, the accuracy of the algorithm is independent of IMI inflation because it is based on the combinatorics of 64 binning indexes. The algorithmically estimated conversion probabilities are highly correlated with UCP observed probabilities across multiple cell and nuclei sample types including HEK 293T/NIH 3T3 cells, PBMCs, and mouse lung nuclei, with a MSE of 5.75e-05 (Figure 10). The accuracy of the MCEE algorithm was maintained across a range of sequence depths among these sample types, demonstrating scalability of the approach. This direct comparison against UCP observed values demonstrates highly accurate and flexible estimation of molecule conversion efficiency.

**Figure 10.**
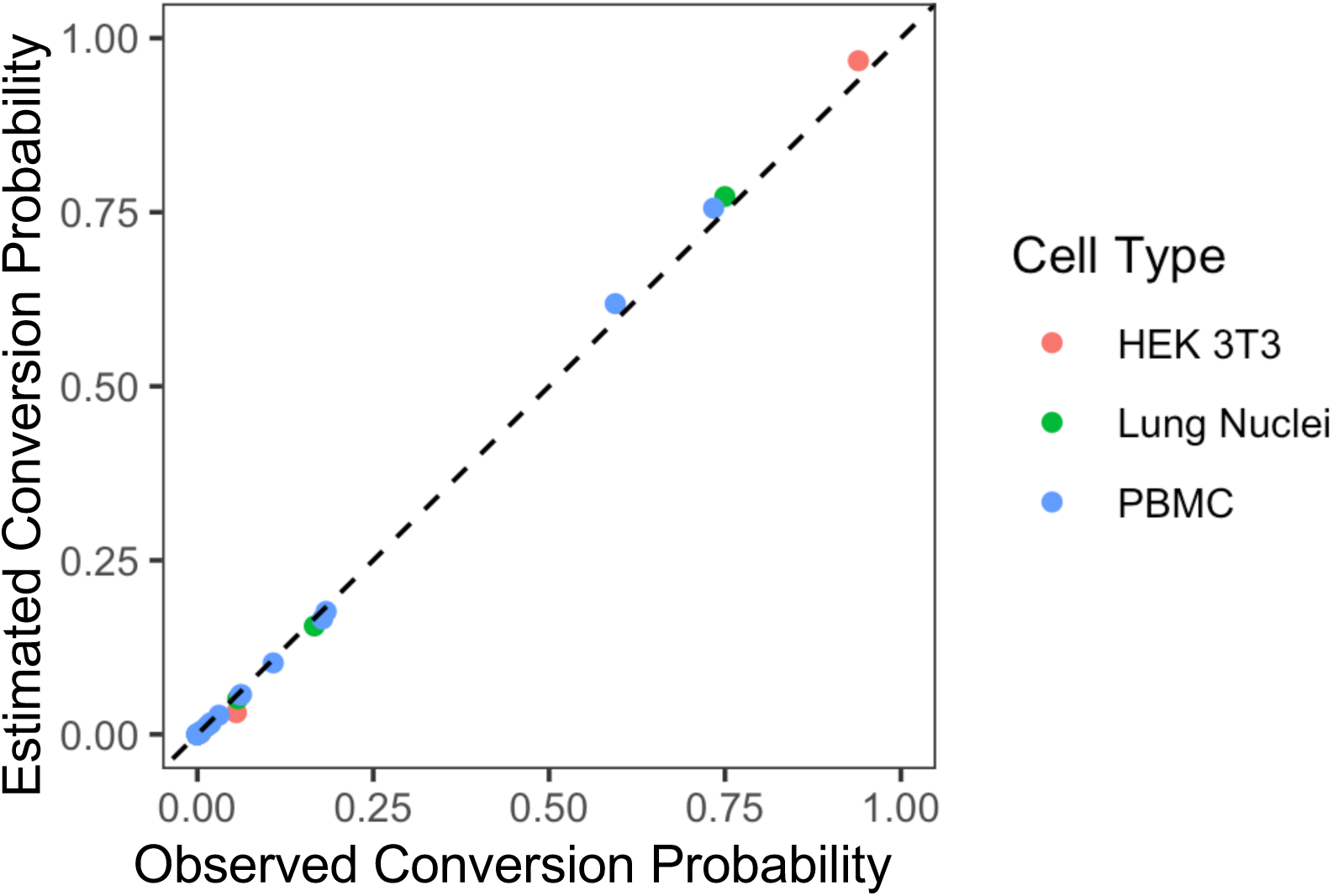
Correlation of IMI inflation probability across three distinct cell types. Correlation of dynamically estimated inflation probability and observed inflation using UCPs in HEK 293T/NIH 3T3, lung nuclei, and PBMCs at varying sequencing depths. Observed probability values come from the true inflated IMI counts associated with UMIs, while estimated values are the result of running DynIMIC using only IMIs from the same dataset. These datasets included 77,636,656 mapped reads for HEK 293T/NIH 3T3, 61,601,586 mapped reads for lung nuclei, and 176,581,519 mapped reads for PBMCs. The mean squared error (MSE) of the inflation estimation is very low (5.75e-05), indicating that the estimation strategy is highly accurate.

The output of the MCEE algorithm, an estimation of molecule conversion efficiency probabilities, is the end result of an iterative process that considers each possible configuration of between one and fifteen IMIs, and for each IMI configuration considers each possible underlying configuration of parent molecules. The total observed count of BGCs with the IMI configuration is proportionally divided among potential parent molecule configurations as the algorithm progresses. From the PBMC UCP experiments described above, we can compare the estimated counts for each pair of IMI and parent molecule configurations considered in the algorithm to the observed counts of the analogous pair of IMI and UMI configurations. Across all of these component estimations, we see a high degree of similarity between the estimated and observed counts (Figure 11A). The algorithm is structured such that configurations with the same number of IMIs and the same number of parent molecules are considered as a group, and estimates are based on the results of the previous group where the number of parent molecules was increased by one. Aggregating these counts within each number of IMIs from configurations with the same number of molecules demonstrates the combined accuracy within these sub-groups within the algorithm (Figure 11B). For each number of IMIs, the estimated distribution of BGC counts with each potential number of parent molecules clearly replicates the observed count distribution of the number of UMIs associated with the same BGCs.

**Figure 11.**
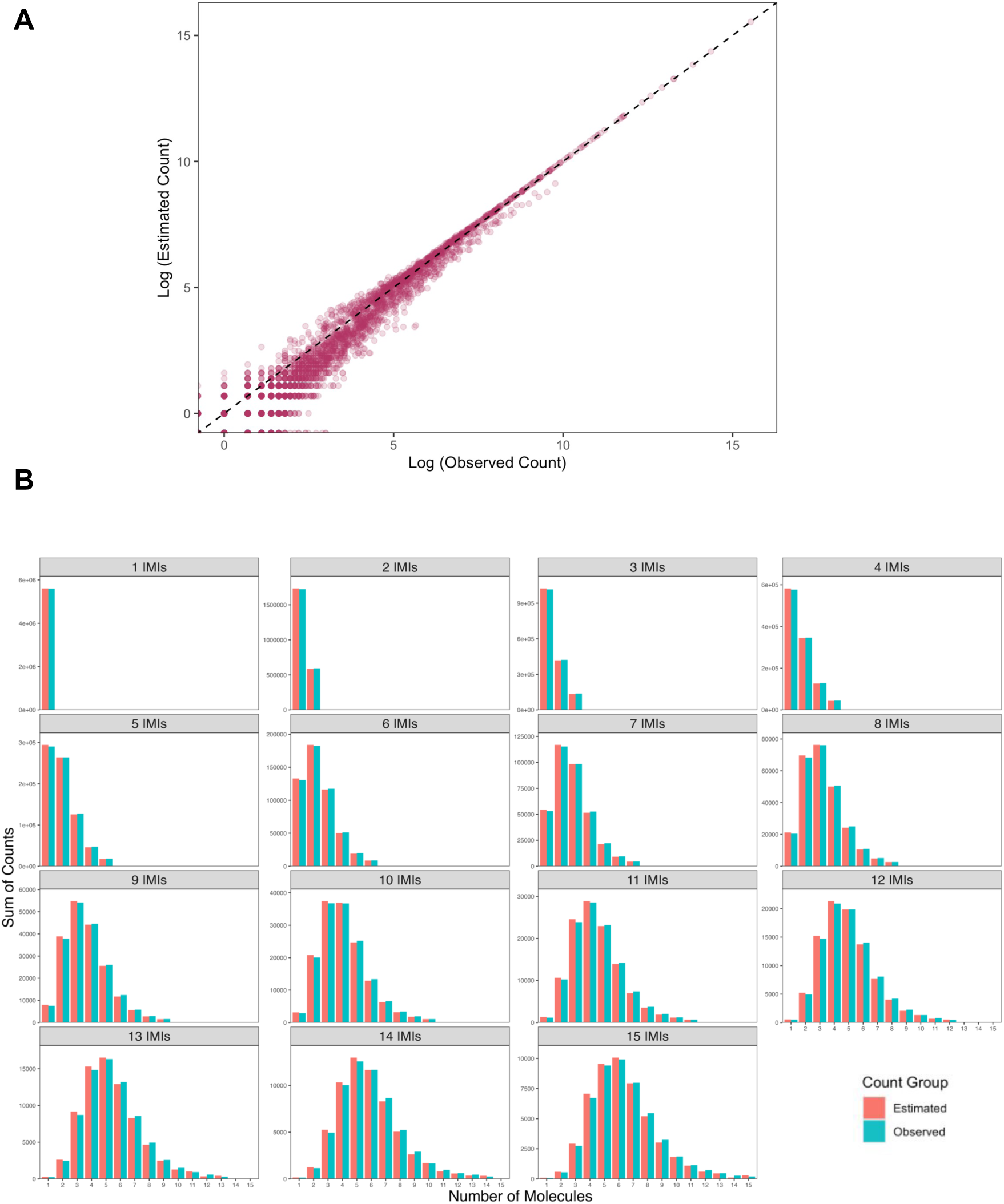
Molecule conversion efficiency estimation accuracy. PBMCs processed through UMI and IMI based analysis from PIPseq V workflow with UCPs. A) estimated counts compared to observed counts for each combination of molecule & IMI configurations evaluated by the dynamic correction algorithm. Observed counts are from UMIs and IMIs with the same configurations as each evaluated molecule & IMI configuration combination. Algorithmic estimation of the proportion of observations from each IMI configuration attributed to each parent molecule configuration is highly concordant with observed counts. B) Sum of observed and estimated counts for each combination of the number of IMIs and molecules from A. For each number of IMIs, the estimated count of observations from each number of potential parent molecules is highly similar to the observed count from UMIs.

### Analytical validation of IMI molecule counts against the UMI-standard

To validate the dynamic IMI correction strategy, we used UCPs to perform UMI- and IMI-based counting across four different sample types: PBMCs, HEK 293T/NIH 3T3, as well as unfixed and fixed mouse brain nuclei, at a wide range of sequencing depths. Relative to the UMI-based approach there is minimal difference in detected sensitivity with the dynamic correction approach (Table 2). In these experiments, the same sequencing data is processed through different analytical pipelines, and therefore the experiments are controlled to a common biochemistry and PIPseq workflow. As anticipated, median genes-per-cell metrics are very well maintained comparing IMI and UMI-based analyses, with a small increase in observed genes noted in IMI-based analysis (1.3% - 6%). We hypothesize that this noted difference in gene mapping arises from small variations in read deduplication and mapping processes in the two pipelines. We also note a relative decrease in detected transcripts per cell in IMI relative to UMI analyses (6% - 13%). These variations may arise from over-correction residual from dynamic IMI correction, but may also reflect stochastic errors native to both UMI and IMI-based counting systems, including sequencing error, and random duplication of UMI and IMI-based identifiers. Identification and evaluation of these analytical process variations will be the subject of future investigation.

**Table 2.** PIPseq V 3’ Single Cell RNA Kits with dynamically-corrected molecular counts in a range of sample types and kit scales produces equivalent sensitivity metrics to UMI-based analysis.

| Sample Type | Kit Scale | Sequencing Depth (Reads Per Cell) | Median Transcripts Per Cell (UMI) | Median Transcripts Per Cell (IMI) | Median Genes Per Cell (UMI) | Median Genes Per Cell (IMI) |
| --- | --- | --- | --- | --- | --- | --- |
| PBMC | T10 | 195,513 | 5,576 | 4,847 (CF = 4) | 2,046 | 2,177 |
| HEK/3T3 | T10 | 31,923 | 11,579 | 10,102 (CF = 2) | 3,649 | 3,700 |
| Mouse Brain Nuclei | T20 | 74,676 | 11,028 | 10,202 (CF = 3) | 3,522 | 3,616 |
| Mouse Lung Nuclei | T20 | 46,686 | 3,080 | 2,869 (CF = 4) | 1,573 | 1,616 |

IMI-based analysis, however, is highly effective at replicating quantitative differential gene expression across diverse cell populations. Heatmaps show the overall expression patterns among cell types for the UMI-based (Figure 12A) and the IMI-based dynamic correction approach (Figure 12B) were very similar, and for each cell type, the top 2000 most variable genes within cell types (Figure 12C) were highly correlated (R≥0.99). These analyses were repeated with mouse brain and lung nuclei preparations with similar demonstrations of concordant differential gene expression (Supplemental Figures) These results demonstrate that the PIPseq V IMI-based approach is effective at both qualitative and quantitative analysis of differential gene expression in complex samples, across multiple cell and tissue types.

**Figure 12.**
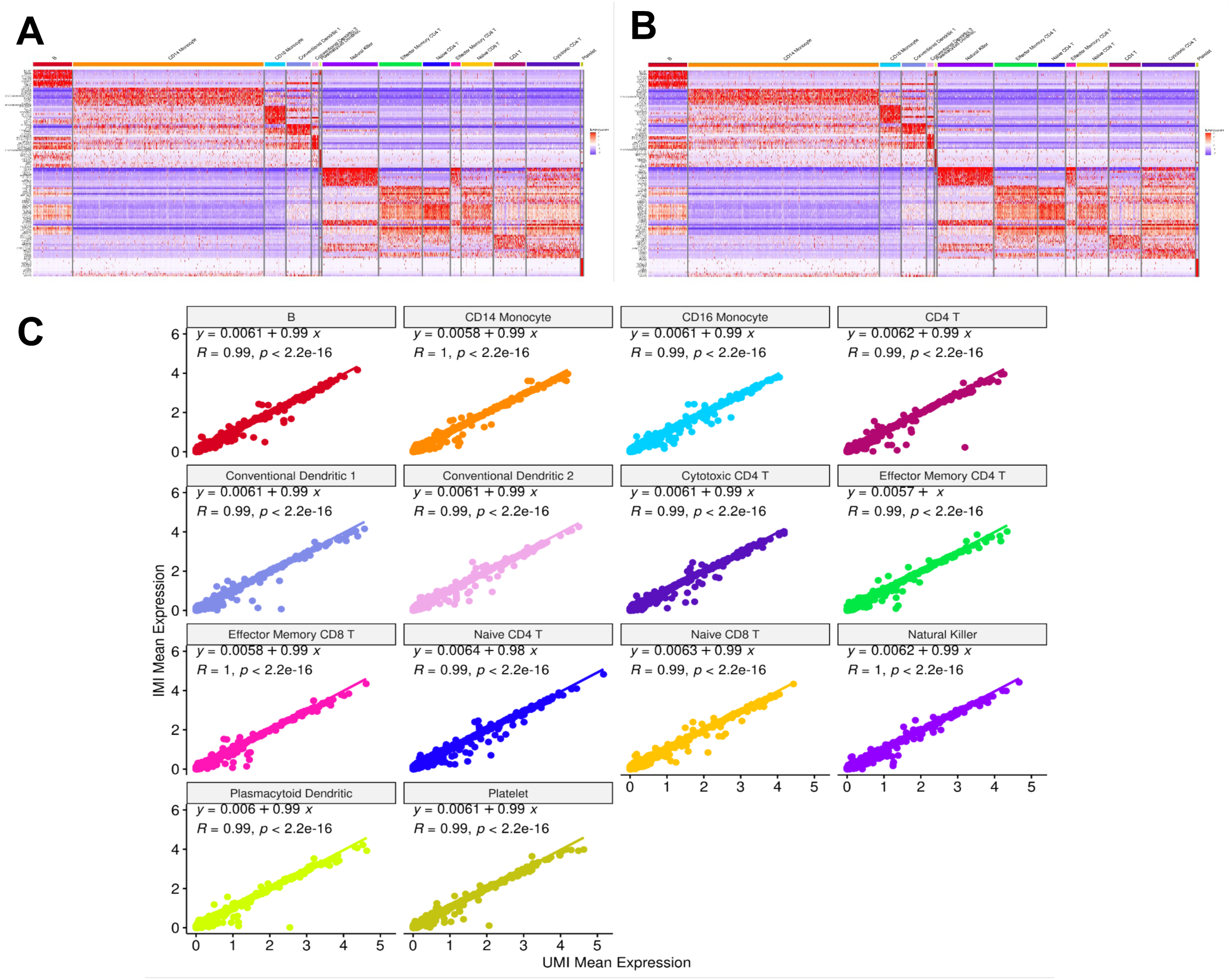
IMI analysis maintains differential gene expression quantitation. PBMCs were processed by PIPseq V using UMI and IMI workflows. A) Differential gene expression heatmap (UMI). B) Differential gene expression heatmap (IMI). C) Correlation of UMI-based and IMI-based quantitative gene expression in individual PBMC cell populations for the top 2000 most variable genes.

We furthermore investigated whether IMI-based analysis vs UMI-based analysis from these samples resulted in observed gene expression bias (Figure 13). The vast majority of genes in each identified PBMC cell population were identified by both IMI and UMI-based analyses (Figure 13A). The majority of differentially expressed genes (abs. log_2_FC > 1.5 adjusted p < 0.01) in PBMC cell populations were shared in IMI and UMI-based analyses based on both the IMI- and UMI-derived count matrices. Of the differentially expressed genes, the genes unique to either IMI- or UMI-based analyses were generally among the least significant. These results further affirm the ability of IMI-based analysis to identify gene expression patterns comparable to UMI-based analysis, and to reliably detect the same differentially expressed genes across cell populations.

**Figure 13.**
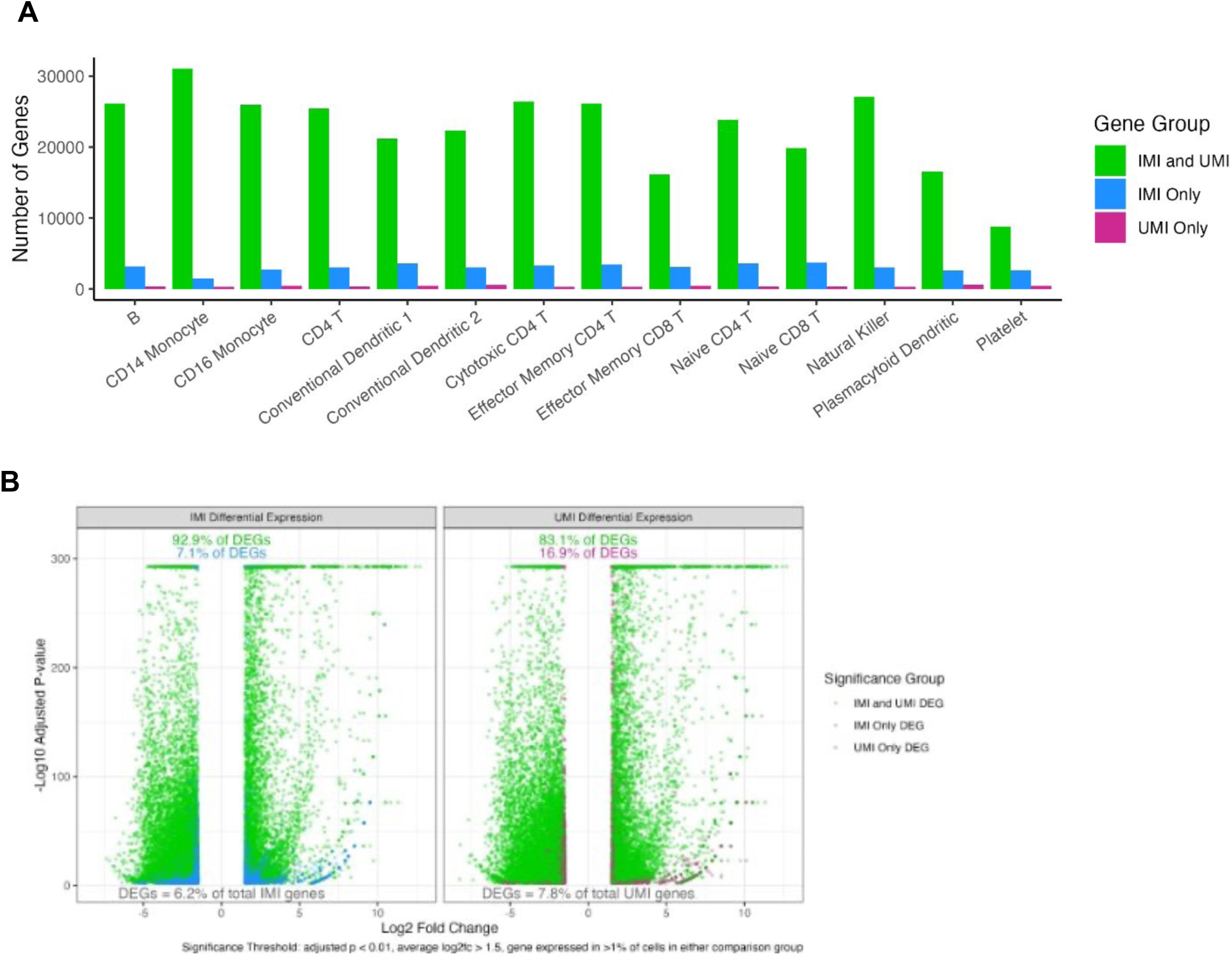
Effect of UMI vs. IMI analysis on observed gene expression in PBMCs. A) Total genes expressed per cell type in UMI and IMI analysis colored by whether the genes were shared in IMI and UMI analyses (green), unique to the IMI analysis (blue), or unique to the UMI analysis (pink). The majority of genes overlap in both analyses. B) Fold changes vs. P-values for differentially expressed genes which represent 6.2% and 7.8% of all genes in the IMI and UMI analyses, respectively. Of the differentially expressed genes, ∼93% are shared among IMI and UMI analyses when using the IMI count matrix. When using the UMI count matrix, 83.1% of all differentially expressed genes are shared between IMI and UMI analyses.

## DISCUSSION

The development of PIPseq V using IMI-based transcript counting represents a significant advance in the field of single-cell genomics. PIPseq V provides high-quality, scalable, flexible, and easily accessible single-cell analysis that can be implemented in any laboratory. Its flexible workflows and sample compatibility have been demonstrated against a diversity of cell and tissue types.

One of the key innovations to the PIPseq workflow is the introduction of intrinsic molecular identifiers (IMIs) for transcript count estimation and elimination of traditional long randomer molecular identifier sequences. This approach has advantages in improved transcript capture specificity and sequencing read utilization, as well as a small observed improvement in gene identification due to mapping efficiency. The combination of IMI-based workflows with improvements to sample processing and mRNA conversion chemistries results in significant improvements in single-cell transcript and gene sensitivities compared to previously released PIPseq chemistries.

A key innovation that enables IMI-based analysis is the introduction of a short, randomer Binning Index that allows quantification of variation in the propagation of randomly fragmented amplicons. This quantification can be solved from only the sequencing data from an individual experiment without additional controls and is robust across variations in sequencing depth, cell type, sample composition, and experimental performance variation. Dynamic correction, performed according to these variations, therefore provides a stable and portable analytical framework that can be applied across single-cell analyses and may have broad applicability beyond single-cell applications as well.

## METHODS

### Human-mouse cell line mixing

Human-mouse mixed cell experiments used cell lines mixed in a 50:50 ratio before cryopreservation. Human HEK 293T cells (ATCC, CRL-3216) were cultured in DMEM (Thermo Fisher, 11995073) with 1% penicillin–streptomycin–glutamine (Thermo Fisher, 10378016) and 10% fetal bovine serum (FBS; Thermo Fisher, A3840001) added as supplements. Mouse NIH 3T3 cells (ATCC, CRL-1658) were cultured in DMEM (Thermo Fisher, 11995073) with 1% penicillin-streptomycin-glutamine and 10% bovine calf serum (ATCC, 30-2030) added as supplements. Once the cells reached 70% confluency, they were harvested and centrifuged at 200xg for 5-minutes centrifugation. Supernatants were removed, and cells were resuspended in 1mL of their respective media and mixed gently with a wide-bore P1000 pipette. Each cell type was counted using a LUNA-FL™ Dual Fluorescence Cell Counter. Cells were dyed with Acridine Orange/Propidium Iodide Stain (Logos Biosystems, F23001) to count the viable nucleated cells. Cells were mixed 1:1 ratio followed by 200xg for 5 min centrifugation. After removing the supernatant, mixed cells were resuspended at 1X10^6^ cells/mL concentration in chilled CryoStor® CS10 media frozen in liquid nitrogen.

### Cultured cell mixture preparation

Frozen aliquots of Human-mouse cell mixture were thawed in a 37°C water bath for 2 minutes or until 70% of the freezing media thawed and transferred to a 15 mL conical tube containing 9 mL pre-warmed cell media (DMEM with 10% fetal bovine serum). Cells were centrifuged 200xg for 5 minutes. Supernatant was removed and the cell pellet was resuspended in 1mL of warm Cell Suspension Buffer (Fluent BioSciences, FB0005067) by gentle mixing. Cells were centrifuged at 200xg for 3 minutes. Supernatant was removed and cells were resuspended into chilled Cell Suspension Buffer and counted using LUNA-FL™ Dual Fluorescence Cell Counter and Acridine Orange/Propidium Iodide Stain. Cell loading concentrations were optimized based on the kit size used.

### Human PBMC preparation

Human Peripheral Blood Mononucleated Cells (PBMCs) cells were sourced from AllCells (Alameda, CA). Cells were thawed in a 37°C water bath until 70% of the freezing media was thawed. Then they were carefully transferred to a 15 mL conical tube containing 9 mL pre-warmed PBMC thawing media (RPMI 1640 Medium (Thermo Fisher, 11875119) and 10% fetal bovine serum). Cells were then centrifuged at 350xg for 5 minutes. Supernatant was removed and the cell pellet was resuspended in 1mL of pre-warmed Cell Suspension Buffer, mixed well, and centrifuged for a second time at 350xg for 3 minutes. PBMC pellets after supernatant removal were finally resuspended using chilled Cell Suspension Buffer, counted, and optimized as described above.

### Nuclei preparation

Nuclei were prepared using the Fluent V Nuclei Isolation Kit (FB0005375) according to the Fluent V Nuclei Isolation Kit User Guide (FB0003716). Cryopreserved frozen brain tissues (CellBiologics, MT118) and lung (CellBiologics, MT130) from mouse strain C57BL/6 were weighed to obtain an appropriate input mass (<50mg). 1 mL of ice-cold Nuclei Extraction Buffer was added to aid the transfer of preweighed tissue into an open sterile dish on ice and minced into 0.5-1 mm pieces using a sterile scalpel blade. Cut tissue pieces were transferred to a WHEATON® dounce homogenizer (DWK Life Sciences, 357538) using a sterile wide-bore P-1000 tip. Tissue pieces were mechanically dissociated using a loose pestle followed by a tight pestle. Tissue slurry was filtered through a 40 µM pluriStrainer® filter (Pluriselect, 43-50040-51) and collected into an appropriately sized conical tube on ice. The filter was washed with an additional 3 mL of Nuclei Extraction Buffer to collect any residual nuclei trapped in the filter. Filtrate was incubated on ice for 5 mins, and centrifuged at 4 °C in a swinging-bucket centrifuge at 500xg for 5 min. The supernatant was discarded and the pellet was resuspended in 5 mL of 1X nuclei suspension buffer to wash any residual impurities and centrifuged at 500xg for 5 min at 4 °C to collect the nuclei. The supernatant was discarded and for unfixed nuclei samples the nuclei pellet was resuspended at a concentration of 10000 nuclei/µL for T100 in 1X Nuclei Suspension Buffer with BSA and RNase inhibitor. For fixed samples, 1 million nuclei were resuspended in 200 µL 1X Nuclei Suspension Buffer and 820 µL chilled fixative was added as per the Fluent DSP-Methanol Fixation for Nuclei Demonstrated Protocol (FB0004745). After a 15 minute incubation the fixative was quenched with 20.4 µL 1 M Tris pH 7.5. The fixed nuclei were then topped with 2mL of 1X Nuclei Suspension Buffer, centrifuged at 500xg for 5 min at 4 °C, supernatant was discarded and nuclei pellet was resuspended in 1X Nuclei Suspension Buffer without BSA or RNase inhibitor at a concentration of 5,000 nuclei/µL for T20.

### PIPseq workflow

The PIPseq™ V Single Cell RNA Kit utilizes a standardized workflow unless stated otherwise, spanning 4 different cell input sizes with different reagent input volumes at each step. For comparison experiments that used v4PLUS workflow and chemistry, refer to the PIPseq™ v4PLUS User Guides (FB0001026, FB0002130, FB0003657). (For 5,000 cell input, refer to PIPseq™ V T2 3ʹ Single Cell RNA Kit User Guide (FB0005260). For 17,000 cell input, refer to PIPseq™ V T10 3ʹ Single Cell RNA Kit User Guide (FB0004762). For 40,000 cell input, refer to PIPseq™ V T20 3ʹ Single Cell RNA Kit User Guide (FB0005261). For 200,000 cell input, refer to PIPseq™ V T100 3ʹ Single Cell RNA Kit User Guide (FB0005262).

### PIPseq core-template particle configuration

PIPseq bead synthesis has been described previously ^11^. The PIPseq V barcoding oligonucleotide comprises a PCR handle, cell barcodes, a 3-base binning index, a fixed spacer sequence, and a poly(dT)V capture site for 3′ capture of mRNA. For comparative experiments, UMI-containing particles (UCPs) replacing the binning index and spacer sequence with a traditional N12 UMI were also synthesized (Figure 3).

### DNA sequencing

Libraries were pooled with 1-2% PhiX and sequenced using Illumina NextSeq 2000 (P3 200 cycles), NovaSeq 6000 (S4 Reagent Kit v1.5, 300 cycles), or NovaSeq X (25B Flow Cell, 300 cycles) instruments. PIPseq employs different sequencing read structures depending on the library type. v4.0PLUS and UCP libraries utilize a 54-base Read 1 (R1), followed by optional 8 or 10 base indexes (I1/I2) for sample demultiplexing. A minimum 63-base Read 2 (R2) is required for these libraries. PIPseq V (IMI) libraries differ with a shorter 45-base R1, followed by the same optional index options. However, PIPseq V R2 requires at least 72 bases for analysis. Oligonucleotides used in this study are described in PIPseq V User Guides (see PIPseq Workflow above). PIPseq libraries are designed to fit Illumina 100-cycle reagent cartridges for cost-efficient sequencing.

### Analytical Methods - The molecule conversion efficiency estimation algorithm

The MCEE algorithm (Figure 4) estimates conversion efficiency from a single parent mRNA transcript (molecule) to cDNA copies, which are randomly fragmented to produce IMIs. This estimation is possible because the maximum number of IMIs from a parent molecule for a certain number of amplification cycles is known, and because of the presence of the randomer binning index, which is shared by IMIs originating from the same parent molecule.

The binning index consists of 3 bases, so there are 64 (*4^3^*) possible bins. For any molecule, the associated BI can be thought of as the outcome of a random selection from the 64 possibilities. For a set of molecules, the probabilities of having certain selections of BIs can be calculated using combinatorics. For example, consider a set of three molecules A, B, and C. There are 64^3^ = 262, 144 possible ways to assign BIs to each of these molecules, since there are 64 possible BIs which can be selected for each of the molecules. The chance of having any specific selection, such as: molecule A has BI 2, B has BI 36, and C has BI 64, is therefore 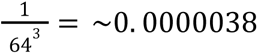. If we are only interested in the probability of selecting the BIs 2, 36, and 64, there are 3 * 2 * 1 = 6 ways to assign the BIs to molecules A, B, and C, so the probability of having this combination of BIs is 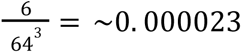. This probability will be the same for any combination of three distinct BIs. Considering all the selections where the three molecules will have different BIs gives 64 * 63 * 62 = 249, 984 ways for this to occur, so the probability is 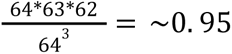. The sum of the selections where the three molecules have different BIs (249, 984), the selections where there two of the molecules share a BI (64 * 63 * *C*(3, 2) = 12, 096) and the selections where the molecules all have the same BI (64) sum to the total 64^3^ possible BI assignments. Within the MCEE algorithm, we consider molecule configurations under this framework, where the BIs themselves are not important. The probability of getting a certain configuration of *m* molecules is the probability of selecting with replacement *m* times from 64 possible binning indexes, and getting a selection equivalent to the given configuration.

Calculating the probability of different distributions of binning indexes among molecules is simple because the number of binning indexes is known. However, the probabilities associated with distributions of binning indexes among IMIs are more complicated, because two IMIs with the same binning index can either be a result of inflation, where they are expected to have the same binning index since they share the same parent molecule, or they are IMIs from distinct molecules, in which case the probability of sharing a binning index comes from the combinatorics of selecting from the 64 possible BIs. In a dataset with no IMI inflation, the probability of getting a certain IMI configuration among BIs would be equivalent to the same configuration of molecules. With greater conversion efficiency, and greater inflation, these probabilities shift. Estimating the conversion efficiency requires inferring the relative composition of parent molecule configurations underlying the observed IMI configurations. This is made possible by limiting the WTA cycles so that the maximum number of IMIs which can share a parent molecule is known. Therefore, for each distribution of IMIs among bins, all possible parent molecule configurations are known. At most, each IMI represents a unique molecule. The known cycle number means that the minimum possible number of parent molecules in each bin in a configuration is also known, it is the result of ceiling dividing the number of IMIs in the bin by the maximum possible number of IMIs from one molecule. Additionally, there are some configurations of IMIs which have only one possible parent molecule configuration. This is the starting point for the MCEE algorithm.

Molecule conversion efficiency estimation is performed using the observed counts for each possible configuration of IMIs in bins, from 1 IMI to the maximum possible number of IMIs that can be created from one parent molecule (15 for 5 WTA cycles). Within the given dataset, the number of observations of equivalent distributions of IMIs in BIs (the same IMI counts in different bins, regardless of the exact BIs) are counted across BGCs with up to 15 total IMIs. Then, for each number of IMIs *i*, and each configuration of IMIs, all potential configurations of parent molecules that can produce the IMI distribution (considering all possible IMI inflation scenarios) are assessed. The starting point for each *i* is the IMI configuration where each IMI is in its own bin, and the number of parent molecules (*m*) is therefore definitive and is equal to the number of IMIs (*i*). This IMI configuration is significant because there is only one possible parent molecule configuration, and only one inflation scenario (there is no IMI inflation).

The total pool of *i* IMIs derived from *m* molecules where *m=i* is estimated by dividing the known observations by the expected fraction of *m* molecules as follows: For each number of molecules *m*, let *T_m_* be the number of instances of uninrflated *m* molecules, and *O_m_* the number observed uninflated *m* molecule in *m* bins. *T_m_* can be derived by:

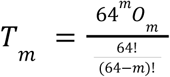

For example:

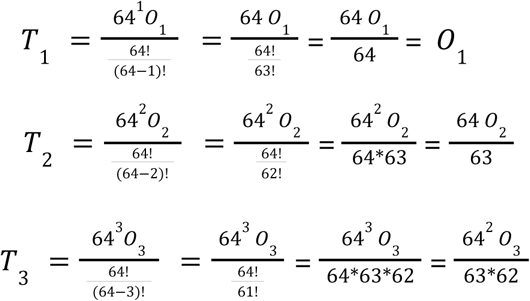

For every other possible configuration of *m* molecules (in fewer than *m* bins), the estimated number of occurrences of the configuration in the dataset is calculated by multiplying *T_m_* by the probability of getting that molecule configuration when *m* molecules are distributed among 64 bins (because we are considering molecules, the probability depends only on BI combinatorics). Then, the expected number of observations of each distribution of *i* IMIs that can be created by this molecule configuration is taken by proportionally splitting the estimated occurrences of the molecule configuration by the probability of getting each IMI configuration from the molecule configuration (product of inflation probabilities from each molecule to each IMI for each configuration of *i* IMIs from this distribution of *m* molecules). The estimated occurrences are subtracted from the remaining observed counts for each IMI distribution (Figure 6C) in order to obtain the estimated (uninflated) IMI counts for each of the 15 IMI distributions. For the first pass where *i* equals *m*, the molecule configuration is equivalent to the IMI configuration, so there is only one possibility. For subsequent values of *m*, there are *i-m* IMIs which are inflated, and the total pool of molecules *T_m_* which has *i-m* inflated IMIs is estimated as before. We consider each *m*, and thus each inflation value *i-m* separately because this allows us to restrict the scope of our probabilities within a certain inflation regime.

Subtracting the estimated occurrences from *m* molecules for each configuration of *i* IMIs allows us to subsequently estimate the occurrences of the same configuration from *m-1* molecules, since the remaining counts for each distribution of *i* IMIs no longer include observations from *i* to *m* molecules. For example, the distribution of *i* IMIs with three IMIs in one bin and two IMIs in another bin (*I_3,2_*) can result from at least two molecules (three IMIs are inflated) and at most five molecules (uninflated). In the first pass, where *i=5* and *m=5*, the estimated occurrences of *I_3,2_* from *m* molecules are calculated by obtaining *T_5_* from the distribution of five IMIs in five bins (O_5_). *T_5_* is then multiplied by the probability of getting the molecule distribution *M_3,2_*:

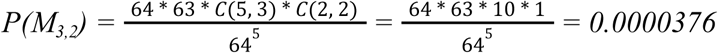

when 5 molecules are distributed among 64 bins. Since the molecules are uninflated, the estimated occurrences of *I3,2* from five molecules are given. This estimate is then subtracted from the observed counts of *I_3,2_*. In the next pass, where *i=5* and *m=4*, *T_4_* with one inflated IMI is estimated by using the remaining counts of the IMI distribution *I_2,1,1,1_*with five IMIs in four bins (O_4_). The remaining counts for *I_2,1,1,1_*now only contain the estimated occurrences from four molecules, since estimated occurrences from five molecules were subtracted in the previous pass. The possible molecule distributions that can produce *I_3,2_*include M_3,1_ and M_2,2_. The estimated occurrences of each of these molecule distributions are obtained by multiplying *T_4_*by the probability of getting the distribution when four molecules are distributed among 64 bins. Adding an inflated IMI to M_2,2_ will always give the IMI distribution *I_3,2_*, while M_3,1_ will produce *I_3,2_* one in four times (and *I_4,1_* otherwise). The estimated occurrences of M_3,1_ are split proportionally between *I_3,2_*and *I_4,1_.* The total estimated occurrences of *I_3,2_*for *m=4* (the sum of the estimates of *I_3,2_* from M_3,1_ and M_2,2_) is subtracted from the remaining counts of *I_3,2_*. The remaining counts can then be used to estimate the occurrences of *I_3,2_* when *m=3*, and so on. Note that when *m=3,* there are two inflated IMIs, so the conversion probabilities must be included in the estimation. The molecule distribution *M_2,1_* can produce *I_3,2_*either by having two molecules each with one inflated IMI (one in each bin) or by having two inflated IMIs from one molecule in its own bin. The total occurrences of *M_2,1_* from T_3_ with two inflated IMIs are first calculated by multiplying T_3_ by the probability of *M_2,1_*, then splitting this estimate by the relative conversion probabilities (four ways to select one inflated molecule multiplied by the probability of two uninflated molecules and one inflated with two IMIs, and *C(4, 2)=6* ways to select two inflated molecules multiplied by the probability of one uninflated molecule and two molecules with one inflated IMI). The estimates within these two inflation configurations are then further split between the possible resulting IMI distributions in each case (*I_3,2_* and *I_4,1_*), and subtracted from the remaining counts of the IMI distributions. For values of *m* and *i* with more than one inflated IMI, the probabilities of each potential configuration of inflated molecules that could be assigned to *m* molecules is thereby accounted for.

This process of estimating the number of observations for each IMI distribution which can be attributed to each potential parent molecule distribution is performed for each number of molecules m from *m*=*i* down to *m*=1. When *m*=1, the remaining observed count of *i* IMIs in 1 bin becomes the count for *i* IMIs from 1 molecule in the conversion efficiency probability distribution. Because this process starts at *i*=1 and goes up to the maximum possible *i*, for any number of IMIs *i*, the conversion distribution has already been calculated for values from 1 to *i*-1, giving all the probabilities necessary to estimate the counts from each molecule distribution for each distribution of *i* IMIs. If the remaining count when *m*=1 is 0 at any point (there are no instances of *i* IMIs from a single molecule), the MCEE algorithm terminates.

## Acknowledgments

This work was generously supported by grants 1 R44 GM145185-01 and 1R44 GM137648 from the National Institute of General Medical Sciences (NIH). Sequencing performed at the Yale Center for Genome Analysis is supported by NIH grant 1S10OD028669-01.

## Author Contributions

Conceptualization, K.M.F., S.K., R.H.M; methodology, K.M.F., Y.A., S.B., A.A.M., Y.X.; investigation, Y.A., S.B., A.A.M., C.H., Y.X., J.S.A.I., R.K., L.Y., P.H., H.D.R., J.Q.Z., A.O., C.D. T.S., S.C., M.R., K.S.M.; formal analysis; Y.A., S.B., A.A.M, C.H.; paper preparation; K.M.F., Y.A., S.B., A.A.M., Y.X., J.I., P.H., H.R., J.Z., C.D. T.S., S.C., M.R., K.S.M., R.H.M.; Supervision K.M.F., K.S.M., S.K., R.H.M.

## Data Availability

PIPseq datasets used for this publication are available upon request. Additional PIPseq data is available for evaluation at https://www.fluentbio.com/datasets/.

## Software Availability

PIPseeker software is freely available for download at https://www.fluentbio.com/resources/pipseeker-downloads/.

## Competing interests

All authors are current or former employees of Fluent BioSciences. Fluent BioSciences is developing and commercializing PIPseq reagent kits and methods for single-cell analysis.

---

**Supplementary Figure 1.**
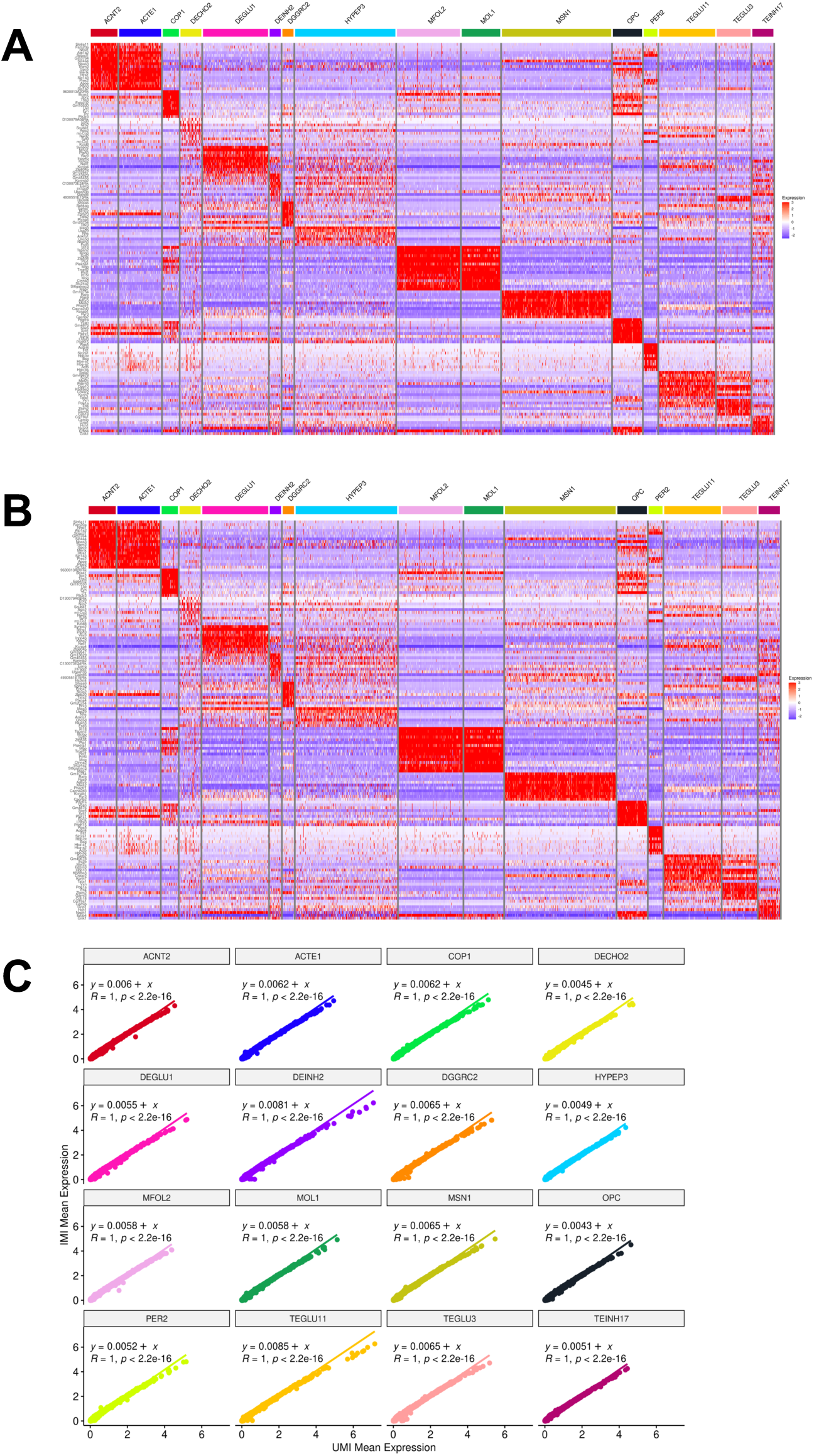
IMI analysis maintains differential gene expression quantitation. Mouse brain nuclei were processed by PIPseq V with UMI and IMI workflows. A) Differential gene expression (UMI). B) Differential gene expression (IMI). C) Correlation of UMI-based and IMI-based quantitative gene expression in individual brain nuclei populations for the top 2000 most variable genes.

**Supplementary Figure 2.**
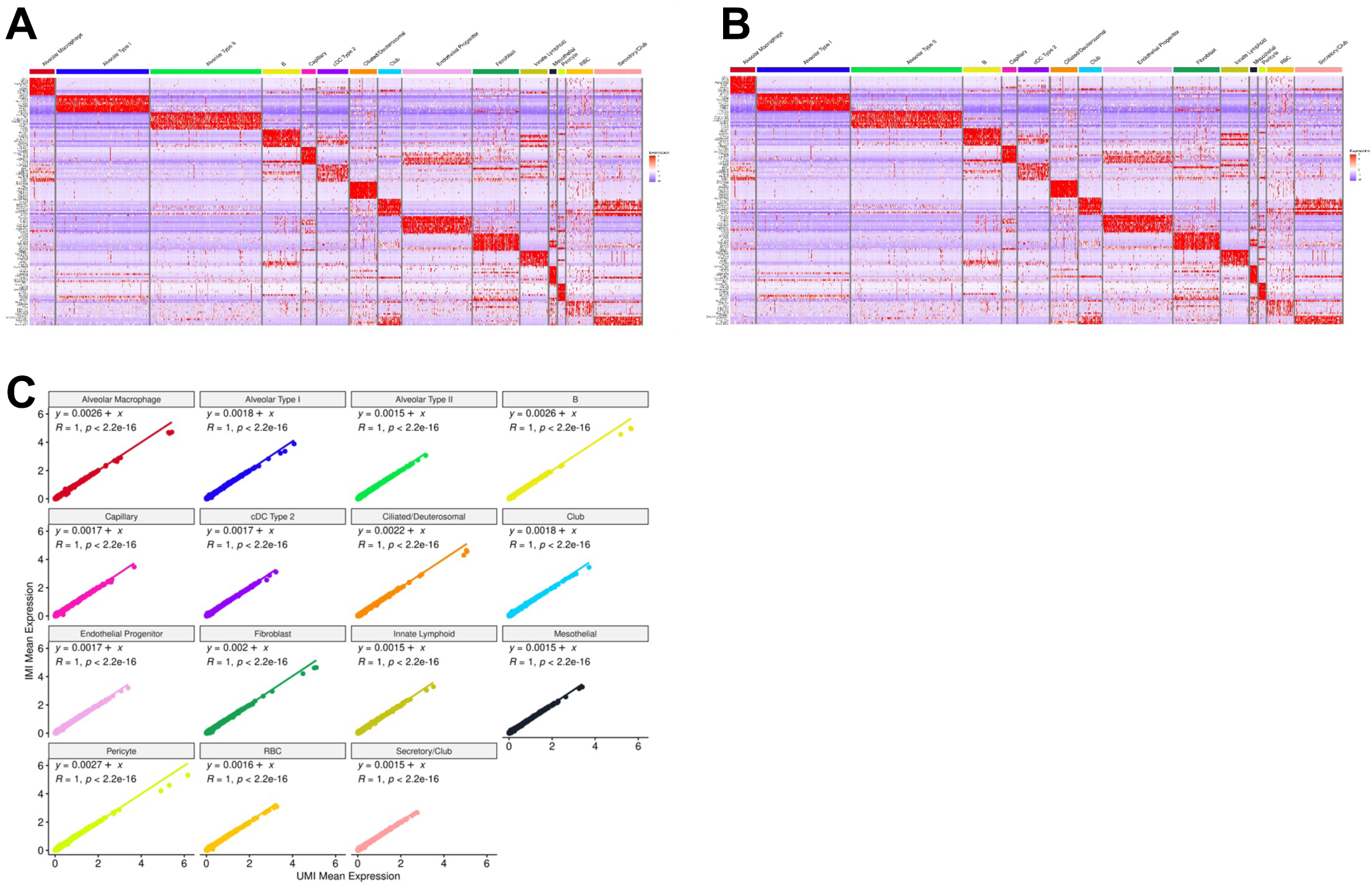
IMI analysis maintains differential gene expression quantitation. Mouse lung nuclei were processed by PIPseq V with UMI and IMI workflows. A) Differential gene expression (UMI). B) Differential gene expression (IMI). C) Correlation of UMI-based and IMI-based quantitative gene expression in individual lung nuclei populations for the top 2000 most variable genes.

